# Disturbance-generated competitive coexistence

**DOI:** 10.1101/2023.11.02.565332

**Authors:** U. A. Trigos-Raczkowski, R. Lyons, M. G. Delgadino, A. S. Ackleh, A. Ostling

**Affiliations:** Department of Integrative Biology, University of Texas at Austin; Department of Mathematics and Computer Science, Karlstad University, Karlstad, Sweden; Department of Mathematics, University of Texas at Austin, Austin; Department of Mathematics, University of Louisiana at Lafayette; Department of Integrative Biology and the Oden Institute, University of Texas at Austin

**Author notes:** A.O designed research; U.T.R, R.L, M.G.D., A.S.A. and A.O. performed research; R.L., A.S.A contributed numerical code; and U.T.R. and A.O. wrote the paper.

**Keywords:** Competitive Coexistence, Population Modeling, Disturbance, Succession, Life history trade-offs

## Abstract

Explaining how competing species coexist remains a challenge in ecology. A major hypothesis is that disturbance opens up the opportunity for types with different “life history” strategies to coexist, allowing types better at getting to and using recently disturbed patches to coexist with better competitor types. A simple model introduced several decades ago demonstrated this, but its focus on patch dynamics (i.e. the dynamics of the number of patches a species occupies) gives limited insight into how coexistence-enabling variation arises from within-patch demographic strategies. Here we present, and demonstrate how to analyze, a partial differential equation model that captures the emergence of larger-scale competitive dynamics from within-patch population dynamics of species competing for patches subject to disturbance. We analyze key cases of the model framework, with competition acting in turn on each aspect of within-patch demography included in the model: reproduction, offspring-survival, and adult-survival. Insights arising from these analyses include: 1) variation between species on a simple reproduction-adult-survival trade-off can enable disturbance-generated coexistence, 2) variation along trade-offs with species’ robustness-to-competition can also generate coexistence 3) disturbance-generated coexistence may or may not involve classical “successional dynamics” within patches, and 4) coexistence is easier to generate at intermediate disturbance rates. Our work here provides new tools for more complete development of the theory of disturbance-generated coexistence.

## Introduction

The contrast between observations of a diversity of coexisting species or types and the intuitive expectation of competitive exclusion favoring the better competitor has been a long-standing topic of discussion for ecologists [1–7]. Disturbance events that reset competitive dynamics are a common theme in many biological systems [8, 9], from forests and coral reefs [10], where storms open up gaps to be newly colonized, to rocky intertidal communities [11], where the action of waves opens up new bare spots on rocks, to pathogen strain communities [12, 13], for which host immune response or death and replacement of hosts can create new patches. The observation of (in some cases) high diversity in these systems, and in particular coexistence of types with different “life history” strategies (different lifetime patterns of reproduction and survival), has led to the hypothesis that disturbance could be a crucial force opening up coexistence opportunities [10, 13–16]. This hypothesis postulates that disturbance creates new patches, allowing types that can better take advantage of the patches to coexist with better competitors who would eventually exclude them.

This hypothesis was formulated in a model focused at the scale of patch dynamics that has since been the subject of a great deal of study [14, 17–21]. The model predicts that variation between species along a “competition-colonization” trade-off can enable their competitive coexistence when disturbance is present. In this trade-off species that are better able to colonize empty patches do indeed coexist with species that are higher up in a presumed competitive hierarchy. Follow-up study of this model advocates for relaxing the strict hierarchy involved for the sake of biological plausibility when considering questions of species packing. This seems to lessen the amount of coexistence possible but does not dismantle this potentially important mechanism [22–24].

Despite the success of this model, its focus on patch-scale dynamics limits the understanding of coexistence it can provide. The model does not incorporate within-patch competitive dynamics, instead presuming the dominant competitor immediately wins in competition for a patch. The trade-off enabling coexistence is defined in terms of a species strategy at the patch-dynamic level. Though the model accounts for variation between species in their ability to get to a new patch, it does not account for variation in their survival nor reproduction in a patch, nor the sensitivity of that demographic performance to competition, which increases in intensity as the patch ages. As a result, some have suggested the model does not capture “successional niche” differences that may also play a role in coexistence when disturbance is present, and are not the same as differences along a competition-colonization trade-off. A new model was presented to illustrate this concept [25]. However, this model was more for illustrative purposes and did not scale up from within-patch dynamics. Forest ecologists have been developing and studying more complex model frameworks, geared specifically for tree species competition, in the hopes of gaining better understanding of this potential “successional niche” differentiation, and have made some important advances [26–32].

Here we study a simple general partial differential equation model that captures how larger-scale competitive dynamics—in the presence of disturbance—emerge from within-patch demography and population dynamics. We demonstrate how to approach the model analytically where possible and numerically otherwise. Our demonstration includes development of invasion analyses for populations structured by patch-age, which require the consideration that an invading species’ distribution across patch ages will change as it invades. We use these approaches to study key cases of the model, with competition acting in turn on each component of demography the model captures, namely on reproduction, offspring-survival, and adult-survival. These analyses yield new insights into the nature of disturbance-generated coexistence.

For example, we find that species differences along a simple demographic trade-off between reproduction and adult-survival can enable disturbance-generated stable coexistence, so long as competition is acting on reproduction and/or adult-survival (and not on offspring-survival alone). We also find coexistence can result from species variation along a variety of trade-offs with their “robustness-to-competition”, meaning how well they maintain aspects of their demographic performance in a patch as species’ densities become high. Another interesting observation we found is that disturbance-generated coexistence can occur with or without within-patch dynamics that involve replacement of one species by another (i.e. classical “successional dynamics”). This is because if density-dependence on recruitment is not extreme, dispersal can maintain high within-patch abundance of a species that coexists on the landscape, even if that species would be excluded in an isolated patch. Finally, intermediate disturbance rates allow a species to coexist with a wider range of other life history strategies.

These insights help us understand how life history diversity and within-patch dynamics differ across systems that vary in disturbance rate or in the component of demographic performance most strongly influenced by competition. They, as well as our model framework and demonstration of approaches to its analysis, provide a starting point for further development of the theory of disturbance-generated coexistence.

## Models & Methods

Next we provide the general framework of our model followed by a description of the specific model cases we examine and analytical methods we applied to each case. These cases differ in which component of demography is impacted by competition, i.e. which component of demography is modeled as having “negative density-dependence”: reproduction, offspring-survival, or adult-survival. The analytical methods we demonstrate include deriving the full structure of coexisting species across patch-ages at equilibrium. This allows consideration of the conditions for feasibility of that equilibrium coexistence, as well as analysis of the local stability of such structured equilibria. Our analytical methods also include derivation of dynamical equations for the average abundance of each species through integration of dynamical equations over patch-age. We in particular apply this approach to derive invasion growth rates, and to consider when one species can invade an equilibrium population of the other—a key criterion for “stable coexistence” of species used in ecology [4]. After describing the general framework, and each model case and the analytical methods employed for each, we describe our numerical approach, which we used to validate the analytical results, as well as to extend the analyses to cases not analytically tractable.

### Patch-Age Structured Model Framework

The first component of our model framework is a set of equations capturing the basic disturbance and aging of patches shown in Fig. 1. They are an adaptation of the McKendrick–Von Foerster equation [33], and analogous to Levin and Paine’s model of patch dynamics [34, 35]:

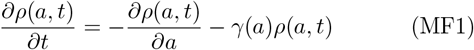

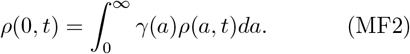

In these equations, *ρ*(*a, t*)da is the proportion of patches with age since disturbance in [*a, a*+*da*] at time t and *γ*(*a*) is the rate at which patches of age a are being disturbed. They together describe the aging of patches, and how disturbance removes patches from an age class and resets them to age *a* = 0. For simplicity, for all of the model cases we analyze here, we take *γ*(*a*) to be a constant rate *γ*, in which case the equilibrium distribution of patch-ages 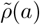 takes the form

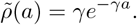

In this case many patches are young and very old patches are rare. See Supplement Sec. 1 for a demonstration that this equilibrium is stable.

**Figure 1:**
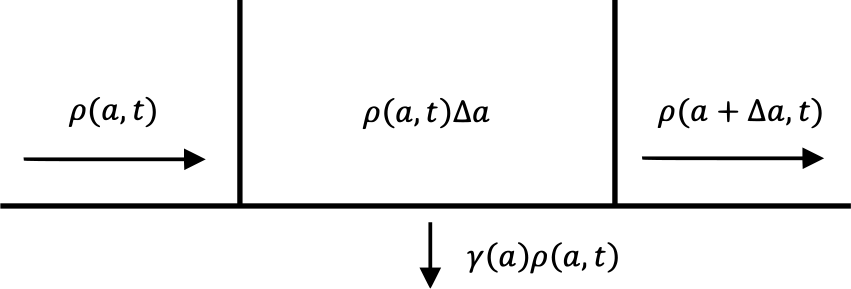
Patch-age structure dynamics. Shown are the flows of patches into and out of a patch-age class between (*a*) and (*a* + Δ*a*). The proportion of patches in that class at time *t* is *ρ*(*a, t*)Δ*a*. Aging causes patches to flow into and out of this age class, and disturbance occurring at a per-patch rate *γ*(*a*) removes patches from the age class, resetting them to age *a* = 0. Consideration of these flows gives the equation 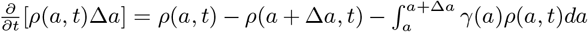 by Δ*a* and taking the limit as Δ*a* → 0 results in (MF1). The resetting of patches to age 0 after being disturbed is accounted for in boundary condition (MF2).

The second component of our model framework describes the dynamics of *n*_*i*_(*a, t*), the abundance of species *i* in a ‘typical’ patch of age a since disturbance. We do not explicitly include stochasticity or other sources of variation across patches in our model, and hence only the age a of a patch distinguishes it. Abundance can be measured in a variety of ways, including number of individuals and biomass. It is often considered per-unit-area and hence referred to as “density”, which we will do here. Note a useful quantity for assessing coexistence is the average density of species i on the landscape, across all patches of all ages, *N*_*i*_, defined as the integral of *n*_*i*_(*a, t*) weighted by the proportion of patches in each age *ρ*(*a, t*), i.e.:

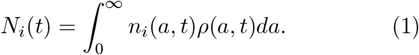

Also note we take *n*_*i*_(*a, t*) to represent the density of *adults* of species i in a typical patch of age a since disturbance, and *N*_*i*_(*t*) to be the average density of *adults* across all patches. The model does not explicitly keep track of the abundance of individuals that have not yet reached the adult stage, i.e. the juveniles, though it factor in aspects of life history before adulthood–in particular their survival to become adults.

The following general model framework encapsulates how the dynamics of *n*_*i*_(*a*) are shaped by the basic demographic processes of recruitment and mortality—which govern all populations—as well as by patch-aging, and specifies the state of the species in newly disturbed patches through a boundary equation:

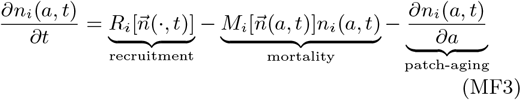

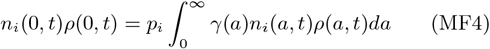

The exact form of (MF3) differs over the model cases we consider and for (MF4) we (in all cases) use a specific form described below.

The first term on the right hand side of (MF3), 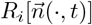, is species *i*’s recruitment of offspring into the patch of age a. It may depend on the abundance or densities of all species—represented by the vector 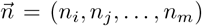—in the patch of age a, as well as in patches of of any other age (indicated by the ““in 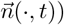. In analyzing our specific model cases we consider just two species labeled *i* and *j*. The recruitment 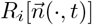 includes two key processes: reproduction and offspring survival to adulthood. We describe each below. The reproduction part of the recruitment rate 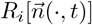 is from all patches from which offspring disperse to the focal patch of age a. Hence it will depend on species’ densities—not just in the focal patch of age a—but across patch-ages, i.e. on the 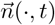. In particular there will be a positive dependence on the focal species density *n*_*i*_(·, *t*) since the more abundant the species is across the landscape, the more it is expected to reproduce. In our model cases we assume equally likely dispersal between all patches, and hence the reproduction part of the recruitment term is simply the average rate of reproduction across patches of all ages on the landscape. If competition is acting on reproduction, the reproduction term will also depend negatively on the density of species *i* and all other species across patch ages, i.e. negatively on 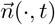. Specifically, the reproduction term for each patch age will depend not only positively on the focal species’ density, but also negatively on the density of all species in the patch.

The recruitment term 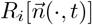 also includes the rate at which offspring survive to adulthood in the target patch of age a. This component of recruitment will only depend on the densities of the species in that patch of age a, i.e. only on 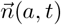, and only if competition is acting on offspring survival, hence the dependence will be negative. We note that our model framework does note explicitly include that it may take time for offspring to become adults.

The next term in (MF3) captures the mortality of adults of species i in the patch, as a per-capita mortality rate 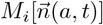, multiplied by the abundance of species i in the patch to calculate the population level mortality rate. The per-capita rate 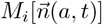 may negatively depend on the abundances of the species in the patch of age a due to competition.

In summary, the recruitment and mortality terms of the model framework can be formulated in a variety of ways to consider the role of competition acting on reproduction, offspring-survival, and/or adult-survival and to consider whether variation between species in each of these aspects of demography and their robustness-to-competition enable disturbance-generated coexistence.

The last term in (MF3) reflects the aging of the patch. The explanation for this term is more subtle than for the aging term in (MF1), because *n*_*i*_(*a, t*) is not a number of things in *a* patch-age class between *a* and *a* + Δ*a*—it instead represents abundance conditional on patch-age. However, the need for the aging term can be seen when considering the density at a time step:

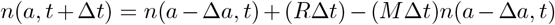

i.e. that the density of a species in a patch of age a at time (*t*+*Δt*) is equal to its value in a patch of age (*a − Δa*) at time *t* (where Δa is the amount of aging that occurred in the time interval *Δt*). Subtracting *n*(*a, t*) from both sides, and dividing by Δt (or equivalently and more usefully Δa = Δt for the term (*n*(*a − Δa, t*) *− n*(*a, t*))) and taking the limit *Δt* → 0 produces an equation including the last term in (MF3).

Finally (MF4) describes the state of species’ abundances in a patch right after it has been disturbed. As written, it allows for a proportion of individuals of a given species *i* (p_*i*_) to survive the disturbance of a patch and be the first to colonize a new patch. One specific mechanism of this would be what is known as “advanced regeneration” in forests [36], in which individuals in the understory survive a disturbance of canopy trees, and are the first to adulthood in the new patch. For simplicity, in all model cases below, we take *p*_*i*_ = 0, giving the simple boundary condition:

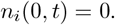

### Competition Acting on Reproduction

The first model case we consider has competition acting only on per-capita reproduction. The more individuals there are in a patch, the less access to resources for reproduction each individual will have, and thus each will produce fewer offspring. Specifically, we consider the following form for model framework (MF3):

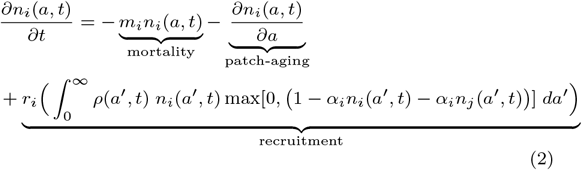

In (2), the integral in the recruitment term gives the average rate of offspring production across patches (those offspring are assumed to disperse uniformly across patches), with the reproduction in each patch impacted by competition, or “negative density-dependence.” That negative density-dependence is modeled as a linearly declining function, constrained by a max function to go no lower than 0, in order to avoid biologically unattainable negative values. The parameter *r*_*i*_ in this recruitment term represents a total reproductive investment. It is equal to the product of species i’s maximal per-capita rate of reproduction (that would occur in new empty patches), times the likelihood of its offspring surviving to adulthood in the focal patch (here independent of the focal densities).

Note however that (2) does not model competition acting on offspring-survival (the recruitment term does *not* depend on the density in the focal patch of age a), nor on adult-survival (the per-capita adult mortality is a density-independent rate m_*i*_). Also note that in this model case (and all others) we take the strength of intra and inter-specific competitive effects on a given species to be equal, i.e. the magnitude of the negative effect of both *n*_*i*_(*a, t*) and n_*j*_(*a, t*) on reproduction of species i is the same competition coefficient *α*_*i*_. We do this to focus on when coexistence is generated by disturbance and succession, since when intra and inter-specific negative effects on a species are equal, coexistence is not possible in the absence of disturbance/succession. (See Supplement Sec. 2.1 for more details.)

Below we describe how the conditions for coexistence can be determined analytically in this model, specifically by deriving criteria for when coexistence is *feasible* (i.e. involves positive abundances) at equilibrium, involves *mutual invasion* (each species being able to invade an equilibrium population of the other species), and is *locally stable* (where we examine small perturbations of species’ populations). We refer the reader to Supplement Sec. 3 for a more detailed analysis of each of these.

### Feasibility of Coexistence Equilibrium

In this model case, the equilibrium density of the adult abundance of species *i* in a patch of age a when it coexists with species *j* is:

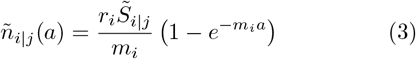

Where 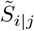 is a constant (has no dependence on a) quantifying the integral portion of the recruitment term in (2) at equilibrium, i.e.

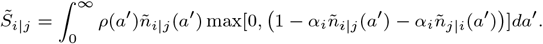

It and the analogous constant for species *j* (coexisting with *i*) can be solved for in terms of model parameters in regions of parameter space where species densities never become so high that they would reduce reproduction to negative values. (See Supplement Sec. 3.3.)

Note that in (3) the term in parentheses is always positive, telling us that the sign of 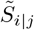 and 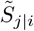 determine whether coexistence is biologically *feasible*, i.e. whether the populations are predicted to have positive abundances. Hence the conditions for two species to have a feasible coexistence equilibrium point in this model case are when

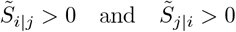

hold simultaneously.

### Mutual Invasion of Average Density Across Patches

To consider model-predicted competitive coexistence as meaningful, the predicted coexistence should be not only feasible (i.e. involve positive population sizes), but also dynamically *stable*, i.e. the system should tend to go towards a state involving coexistence. One definition of “stable coexistence” [4] is when each species can invade a resident population of the other species, i.e. when there is “mutual invasion”. Here we demonstrate such an analysis in the presence of population structure, where it turns out that how an invading population’s structure may change as it invades must be considered. The approach involves integration to get at invasion at the landscape scale, specifically using the average population density.

Our first step was to derive the equilibrium state of each species on its own. In this model case, when only one species is present on the landscape, its equilibrium abundance over patch-ages, which we call *ñ*_*i*|0_(*a*), takes the same general form as *ñ*_*i*|*j*_(*a*) in (3), but the term is replaced with the one species 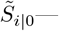 defined as the integral in the recruitment term at equilibrium for species *i* present on its own.

We then derived an analytical expression for the landscape-level invasion growth rate of each species. Specifically, we derived the per-capita growth rate of the average abundance *N*_*i*_(*t*) (defined in (1)) of species *i* across patch ages when started from a very small density, *ñ*_*i*_(*a*) ≈ *ϵ*_*i*_(*a*), where species *j* starts out at its equilibrium density when on its own. The criterion for invasion is for this growth rate to be positive, i.e. for

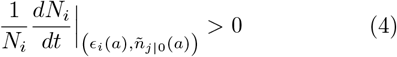

We derived this per-capita growth rate by first considering the dynamical equation for the quantity n_*i*_(*a*)*ρ*(*a*) and then integrating it over all patch-ages. (See Supplement Sec. 3.4 for more details.)

Using this approach, we found that whether invasion is predicted or not depends on the starting structure of the invading species, i.e. on *ϵ*_*i*_(*a*). In particular, we found the invasion could occur for some parameters for which the coexistence is *not* feasible (see Supplemental Sec. 3.4.1). However, we reasoned that the invading species should quickly be distributed over patches in proportion to its equilibrium form (3), i.e.

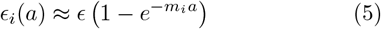

relatively soon after invasion. We found that instead predicting invasion based on this structure results in conditions for mutual invasion that are identical to those for feasible coexistence. (See Supplement Sec. 3.4.2.)

We validated this approach through comparison with numerical simulations (numerical method described below). Specifically, we considered the invasion of a species initially uniformly distributed across patches, i.e. with *ϵ*_*i*_(a) = *ϵ*, and found it does indeed increase in average landscape abundance initially if predicted to analytically under a constant starting density, However, we found that the dependence of its density on patch-age quickly approaches the equilibrium form (i.e. quickly followed (5)), and unless it is also predicted to invade under that starting distribution, the species’ abundance then declines to zero, so that ultimately the invasion is *not* successful (see Fig. 2).

**Figure 2:**
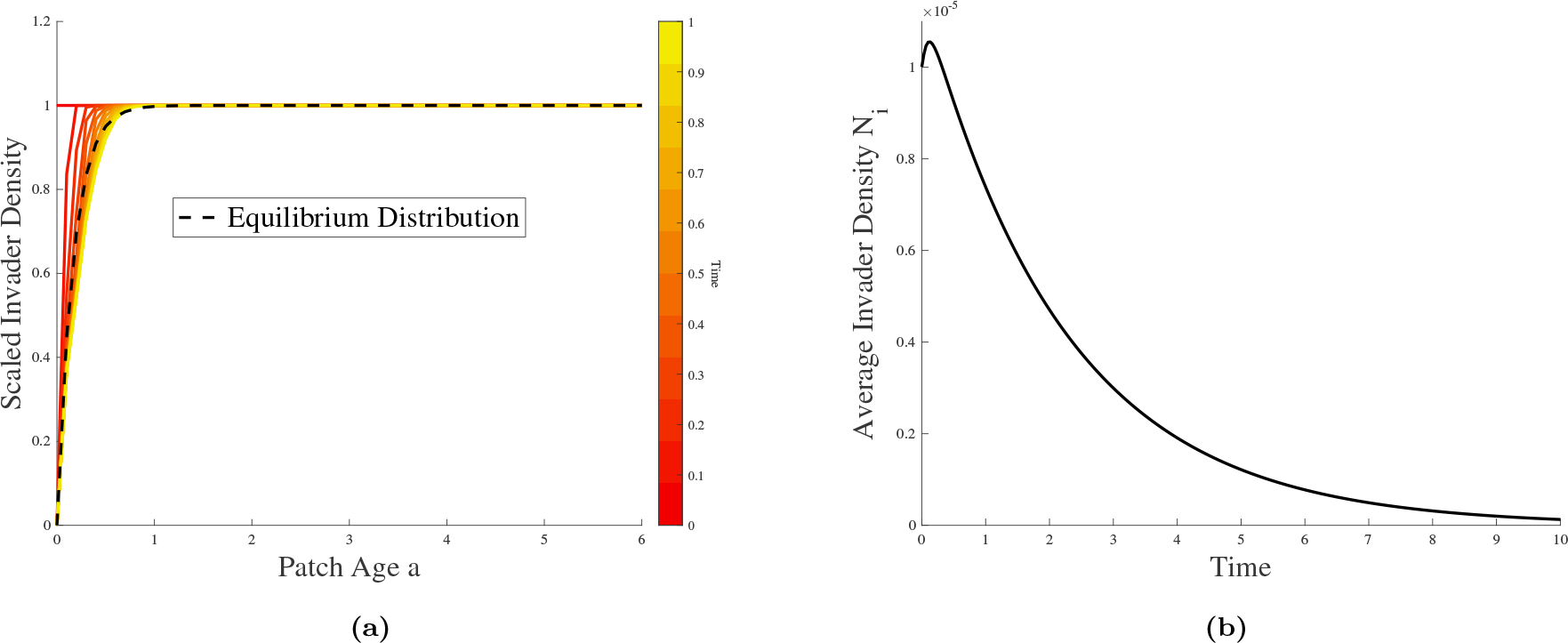
Initial Fast Changes in Invader Population Structure Change the Course of the Invasion. (a) As time increases (red to yellow) the dependence of invader densities on patch-age quickly changes to its equilibrium form (black dashed). Plotted is *n*_*i*_(*a, t*)*/n*_*i*_(20, *t*) for time points *t* indicated by the colors. (b) The average invader density *N*_*i*_ initially increases, but once its structure goes to equilibrium (by *t* = 0.2), it begins to decline to 0 (unsuccessful invasion). (Competition acting on reproduction model case with *γ* = 1 and parameters of the invading species set to *r*_*i*_ = 15, *m*_*i*_ = 6, and of the resident set to *r*_*j*_ = 5.6, *m*_*j*_ = 1.

Hence, the analytical process for determining conditions for *long-term* mutual invasion is to derive the conditions for when (4) and the corresponding condition for species *j* simultaneously hold, with initial abundances *ϵ*_*i*_(*a*) and *ϵ*_*j*_(*a*) assumed to be proportional to that species’ equilibrium structure as in (5).

### Local Stability of Coexistence Equilibrium

We also derived conditions for the local asymptotic stability of the coexistence equilibrium using an approach analogous to standard PDE methods [37, 38] by considering the dynamics of a modified L^2^ norm (ours weighted by the density of patches *ρ*(*a, t*)) of small perturbations *ϵϕ*_*i*_(*a, t*), *ϵϕ*_*j*_(*a, t*) around the coexistence equilibrium densities *ñ*_*i*|*j*_(*a*), *ñ*_*j*|*i*_(*a*) (Supplement Sec. 3.5.2). Our approach yields sufficient but not necessary conditions for local stability. Numerical exploration of those conditions indicate they overlap with those for feasibility and mutual invasion (which as stated above are equivalent conditions in this model), holding wherever they hold.

### Summary of Coexistence Conditions

For this model case of competition acting on reproduction the conditions for a feasible and locally stable coexistence equilibrium of two species (*i* and *j*)—as well as for mutual long-term invasion—can be summarized as

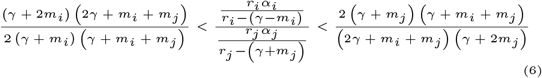

where the right inequality is the condition for feasibility of species *i*/for species *i* to be able to invade species *j*, and the left inequality is the respective conditions for species *j*.

Results for what trade-offs are coexistence-enabling in this model were generated by further analysis of (6) (see Supplemental Sec. 3.6), as well as through numerical exploration of those inequalities. Coexistence results were also validated with the numerical simulation described below.

### Competition Acting on Offspring-Survival

The second model case we consider has competition acting only on offspring-survival. It models the chances of an offspring establishing itself and growing to maturity in a patch of age a to depend negatively on the number of adults already present in a patch of that age. Specifically, we consider the following form of our model framework (MF3):

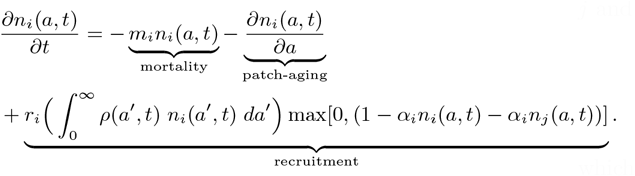

Here the recruitment term is the product of an integral calculating average production of offspring across patches (with *no* negative density-dependence), and a term negatively depending on the density of species in the focal patch of age a, reflecting the influence of competition on offspring-survival to adulthood. That term involves a linear negative density-dependence, limited to nonnegative values by a max function. The rate r_*i*_ again reflects reproductive investment, and should be thought of as the product of per-capita offspring production times maximal offspring-survival (the value it would have in an empty patch). Note the per-capita adult mortality rate m_*i*_ is independent of density.

### Integration to Obtain Dynamical Equations for Average Density

We showed that the average abundance of species governed by this model simply follow the Lotka-Volterra competition model [1, 2] in cases where neither species’ offspring-survival is driven to zero due to competition with adults already established in old patches, i.e. under the assumption that species’ densities remain low enough that they do not cause the linear density-dependent factor of the recruitment term to go to 0 (see Supplement Sec. 4.1). Specifically, we derived a dynamical equation for *n*_*i*_(*a, t*)*ρ*(*a, t*) and integrated it across patch ages to get the following ordinary differential equations for the species average densities *N*_*i*_(*t*), *N*_*j*_(*t*) (defined in (1)):

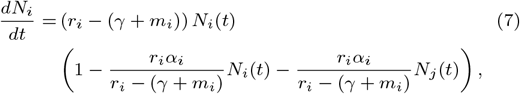

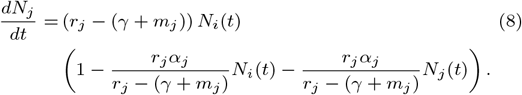

We then used standard approaches for the Lotka-Volterra model to determine coexistence, resulting in the following conditions for species *i* to invade species *j* and for *j* to invade *i* (respectively):

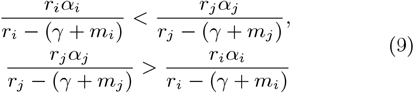

which (clearly) cannot hold simultaneously. Hence in the parameter regime in which the LV equations in (7) and (8) apply, *no coexistence* is possible in this model.

### Invasion Analysis with a Critical Patch-Age

We also derived mutual invasion conditions for this model case that apply *outside* of the regime where offspring-survival is not driven to 0 (see prior section). To do so, we considered a critical patch-age above which an invading species *i* would be blocked, with none of its offspring surviving due to competitive pressure from adults of species *j*. We derived the single species equilibrium structure *ñ*_*j*_(*a*) and then used it to show that a finite critical patch age can only exist when *α*_*i*_ is enough larger than *α*_*j*_ (by a factor depending on other parameters). We then derived the invasion growth rate of the average abundance of species *i* in this regime, using an approach analogous to the prior model case, although here the structure of the invading species does not matter. We then considered when this invasion growth rate for species *i* can be positive, and when species *j*, which is still governed by Lotka-Volterra dynamics because there is no critical patch age for it, can also invade species *i*. This revealed that there *can* be mutual invasion when species i can be blocked from establishing in old patches by species *j*. (See Supplement Sec. 4.2.2 for details.)

### Competition Acting on Adult-Survival

Our final model case considers competition acting on adult-survival, in which the per-capita mortality rate is no longer density-independent, but instead *increases* as species’ densities in patches increase.

This increase in per-capita mortality rate with density translates into a *negative* density-dependence on survival, and on the overall population growth rate in a patch. Mechanisms by which competition might act on adult-survival include less resource availability or easier spread of a pathogen as more adults become established.

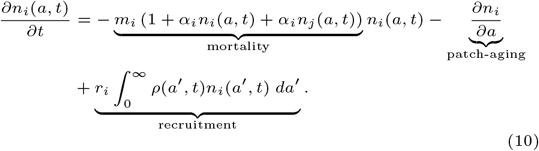

#### Approach

This model case is different from the prior two model cases in that it has a *non-linear* dependence on the density of the focal species in the focal patch of age a. This makes it more difficult to approach analytically. Though we could make some minimal progress with a single species case (See Supplementary Sec. 5), we were unable to solve for integration constants, and hence unable to carry out an invasion analysis analytically. We instead numerically simulated the partial differential equation in (10) using the method described in the next section. We note its results matching with analytical results for the other model cases, providing assurance of accuracy.

### 2^nd^ Order Flux Limiter Numerical Method

The application of numerical schemes to structured models can result in diffusion of the population structure, an inaccurate spreading between age classes [39]. A numerical method used by ecologists to avoid this is the Escalator Boxcar Train [40] which is 1^st^ order accurate. Here instead we use a 2^nd^ order method (code available at [41]) which uses a Flux Limiter approach to limit said numerical diffusion [42, 43]. This method is more accurate than the EBT and has been studied in the context of size-structured population models [43–50]. Numerical experiments suggest this scheme is 2^nd^ order convergent in a smooth setting and the scheme has been shown to behave intuitively in the presence of discontinuities and singularities [42]. (See Supplement Sec. 6 for a detailed description of the numerical scheme.) We use this numerical method both to validate our analytical results (providing assurance of them and the accuracy of this numerical method), and to analyze the competition acting on adult-survival case, where analytical progress was not possible. To calculate regions of long-term mutual invasion, we first simulate one species on its own to equilibrium, and then introduce a small density across patches of the other species. Finally, at a later time, we check if the species’ average density across patches had increased.

## Results

### Coexistence from a Reproduction/ Adult-Survival Trade-Off

As shown in Fig. 3, we found that variation between two species on a reproduction-survival trade-off can lead to disturbance-generated coexistence when competition is acting on either reproduction or on adult-survival. In other words, coexistence can occur between two species *i* and *j* if species *i* is higher in reproduction, *r*_*i*_ > *r*_*j*_, but also higher in mortality *m*_*i*_ > m_*j*_, making species *j* the better survivor, in both model cases. However, in the case of competition acting only on offspring-survival, this trade-off was *not* coexistence generating. (See Table 1 for a summary of our coexistence results by model case.) For the case of competition acting on reproduction, we in particular demonstrated that (6) implies that for any survival advantage of species *j* over species *i*, i.e. any case of higher mortality of species *i*, giving *Δ*_*m*_ = *m*_*i*_ *− m*_*j*_ > 0, there exists a range of reproduction advantages of species *i*, i.e. *Δ*_*r*_ = *r*_*i*_ *− r*_*j*_ *>* 0, that will allow species *i* and *j* to coexist (mutually invade *and* have a feasible coexistence equilibrium that is locally stable). Furthermore, there are no inherent limits to the similarity of the coexisting species. (See Supplemental Sec. 3.6.1.)

**Figure 3:**
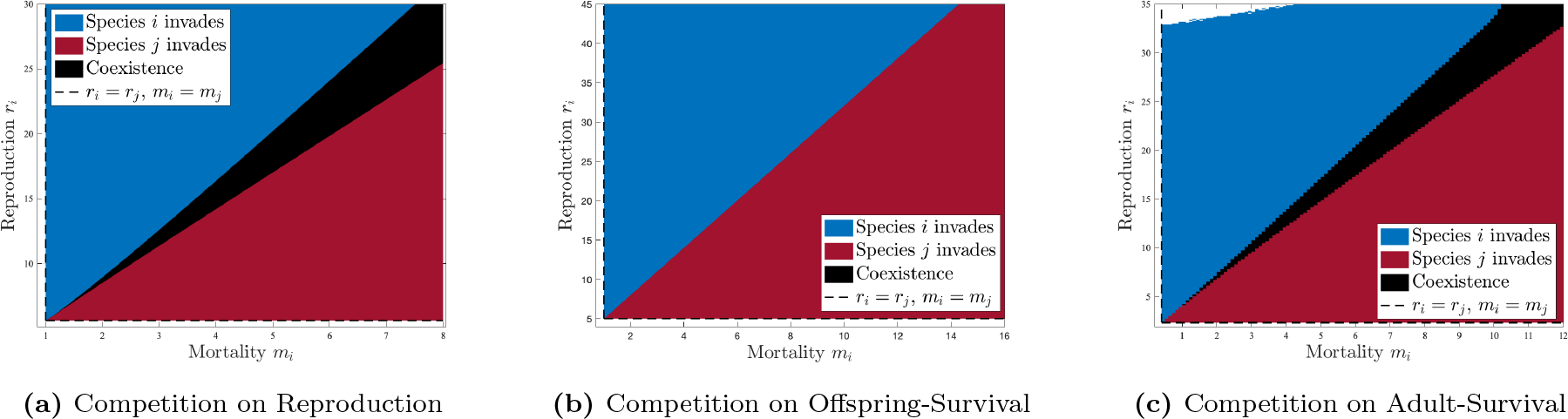
Coexistence from a Reproduction-Adult-Survival Trade-off. Plots all consider potential coexistence between two species (*i* and *j*) when species *i* is higher in reproduction, *r*_*i*_ *> r*_*j*_, but also higher in mortality *m*_*i*_ *> m*_*j*_, making species *j* the better survivor. Each shows for a fixed strategy for species *j*, the outcome of mutual invasion with species *i* within a region defined by the lines *r*_*i*_ = *r*_*j*_ and *m*_*i*_ = *m*_*j*_ (dashed) (species *i* and *j* are equivalent in strategy at their intersection (lower left)). Blue indicates species *i* can invade species *j*, but not vice versa (i.e. species *i* is competitively dominant). Red indicates the reverse. Black indicates the species mutually invade one another and in this sense, species coexistence. When competition is acting on either (3a) reproduction or (3c) adult-survival alone, variation on this trade-off produces coexistence. No such coexistence occurs when competition is acting on (3b) offspring-survival alone. Invasion outcomes were determined using (6) in (3a), using (9) in (3b), and using numerical simulations in (3c). See Supplemental Fig. 1 for verification of (3a) using numerical simulations. Parameters are: (3a) *r*_*i*_ = *r*_*j*_ = 5.6, *m*_*j*_ = 1, *γ* = 0.4 and *α*_*i*_ = *α*_*j*_ = 1.; (3b) *r*_*j*_ = 5, *m*_*j*_ = 1, *γ* = 0.66, and *α*_*j*_ = 0.2.; (3c) *r*_*j*_ = 2.34, *m*_*j*_ = 0.4, *α*_*i*_ = *α*_*j*_ = 1 and *γ* = 1.

### Coexistence from Trade-Offs with Robustness-to-Competition

Each model case yields coexistence from a trade-off involving robustness-to-competition (See Fig. 3 and Supplemental Table 1).

In the case of competition/negative density-dependence on reproduction, the trade-off that generates coexistence is between the robustness of reproduction to competition and adult-survival (see Fig. 3a). If species instead vary on a trade-off between competition robustness and reproduction itself, it will *not* allow coexistence. For this case we demonstrated that for any survival advantage of species i, i.e. lower mortality *Δ*_*m*_ = *m*_*j*_ *− m*_*j*_, there is a range of increased competition sensitivities of species i, i.e. positive *Δ*_*α*_ = *α*_*i*_ *− α*_*j*_ that will allow for coexistence. Furthermore, we showed there are no inherent limits to the similarity of the coexisting species. (See Supplemental Sec. 3.6.2.)

In the case of competition acting on offspring-survival, it is a trade-off between reproduction and robustness of offspring-survival to competition that enables coexistence, i.e. when species *i* has higher reproduction (*r*_*i*_ > *r*_*j*_) but is more sensitive to competition than species *j* (*α*_*i*_ > *α*_*j*_) (see Fig. 3b.) We find this trade-off is only coexistence generating when species i’s sensitivity to competition is high enough that species *j* essentially blocks it from recruiting in older patches. Coexistence through this trade-off does seem to involve an inherent limit to similarity according to our exploration of parameter space so far.

Finally, in the case of competition acting on adult-survival, it is a trade-off between robustness of adult-survival to competition and reproduction that enables coexistence, i.e. if species *i* has higher sensitivity of its adult-survival to competition (*α*_*i*_ > *α*_*j*_), then it can coexist with species j if it also has higher reproduction (*r*_*i*_ > *r*_*j*_) (see Fig. 3c). Our exploration of parameter space using numerical simulations indicated that if the robustness of adult-survival to competition instead trades off with adult-survival itself (i.e. if *α*_*i*_ > *α*_*j*_ and *m*_*i*_ < *m*_*j*_), it would *not* be coexistence generating.

### Coexistence May Not Involve “Successional Dynamics”

The opportunity for coexistence through variation in trade-offs is clearly generated by the incorporation in our model of disturbance resetting patches, and of competitive population dynamics progressing in patches as they age. We constructed our models so that coexistence does not arise without that disturbance and patch-age structure (see Supplemental Sec. 2.1). Thus the within-patch changes of population densities coexisting on the landscape are of particular interest. The classical expectation is for disturbance-generated coexistence to involve species replacing one another in the patch as it ages [10], i.e. “successional dynamics” [51]. In particular, the species investing more in reproduction is expected to be more abundant initially, but to be excluded by a slower reproducing species that can out-compete in other ways as the patch-ages.

In our model cases, the changes in species’ densities as the patch ages among coexisting species can take on a variety of forms that depend on which aspect of demography competition acts on (see Fig. 5). Importantly, when competition acts only on reproduction, coexisting species have their population density simply increase with patch-age to a saturating value and stay there (see Fig. 5a). We showed this holds for all parameter values, in that the monotonically increasing form in (3) always describes the equilibrium dependence of species on patch-age. This result also makes intuitive sense, in that there is no tendency in this model for recruitment into a patch to be more limited as it ages, nor for mortality to become higher. Instead, the production of offspring a species can achieve in a patch wanes as it ages, and hence its contribution to the overall pool of offspring that disperse on the landscape becomes more limited. Note that although the high reproduction species in Fig. 5a does not lessen in abundance in old patches when it is coexisting on the landscape, it *would* get excluded by the high survival species in the absence of disturbance (or in a single patch if it were isolated from the rest of the landscape) (see Supplemental Sec. 2.1).

When competition instead acts on offspring-survival, we recover the classically expected “successional niche” dynamics (Fig. 5b). The high reproduction species is in fact excluded in older patches. Indeed, in this model case, for disturbance to generate coexistence, the high reproduction species has to have a high enough sensitivity to competition to be blocked from recruiting in old patches by its competitor. The only demographic change left to occur in those old patches is mortality, so the eventual exclusion of that species is expected.

Finally, when competition acts on adult-survival, we see patterns of species’ abundance over patch-age in between the other model cases (Fig. 5c). The high reproduction species does achieve higher density early on, reaches a peak value, and then declines in abundance. However, in all the cases we simulated that involved species coexistence, this decline lasts only so long, saturating at a substantial abundance in old patches, where it is balanced by continued recruitment.

### Intermediate Disturbance Yields Larger Coexistence Regimes

Classically in ecology there has been a hypothesis that “intermediate disturbance” will lead to the highest levels of diversity [10], meaning that the highest diversity will occur with disturbances that are “neither very frequent nor infrequent”. Though we are limited in examining this hypothesis through our two-species analyses, we can ask a relevant question, namely how the size of the parameter regime enabling coexistence varies with disturbance rate. Presumably a larger regime of two species coexistence would lead to a larger parameter regime for the possibility of multiple species coexistence as well, and hence a greater tendency for supporting the coexistence of a larger number of species.

Consistently across model cases (and across trade-offs) that can generate coexistence, we found that intermediate values of the disturbance rate *γ* lead to larger coexistence regimes. See Fig. 6 and Supplemental Figs. 2, 4, 6, and 9. When looking at plots of the coexistence regime with a fixed scale (i.e. a fixed range of parameters plotted) while varying disturbance rates, we find the area of the coexistence regime is bigger at intermediate disturbance values. In other words, across the parameter values we examined, for given parameters for species *j*, the range of coexistence-enabling parameters for species *i* within a given rectangular range from species *j*’s parameters tends to be larger at an intermediate disturbance rate (what qualifies as “intermediate” in a quantitative sense varies across different model cases and trade-offs, as well as with species *j*’s parameters).

**Figure 4:**
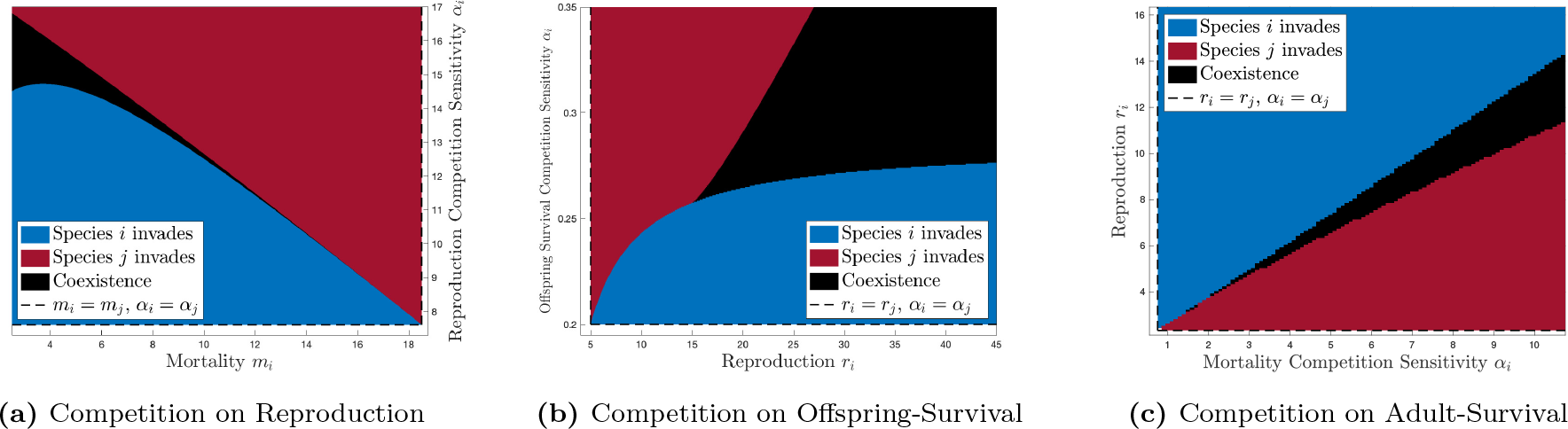
Coexistence from Trade-offs with Robustness-to-Competition. Plots all consider potential coexistence between two species (*i* and *j*) when species *j* is more robust to competition (i.e. *α*_*i*_ *> α*_*j*_), and species *i* is in turn either better at survival (*m*_*i*_ *< m*_*j*_) or better at reproduction (*r*_*i*_ *> r*_*j*_). Each shows for a fixed strategy for species *j*, the outcome of mutual invasion with species *i* within a region defined by lines (dashed) indicating where species *i* is equivalent to species *j* in a given component of demography (with the species entirely equivalent to each other at their intersection). Blue indicates species *i* can invade species *j*, but not vice versa (i.e. species *i* is competitively dominant). Red indicates the reverse. Black indicates the species mutually invade one another and in this sense, species coexistence. Each model case yields coexistence from a trade-off involving robustness-to-competition: (4a) With competition acting on reproduction, variation on a trade-off between robustness of reproduction to competition and adult-survival gives coexistence; (4b) With competition acting on offspring-survival, variation on a trade-off between robustness of offspring-survival to competition and reproduction gives coexistence; (4c) With competition acting on adult-survival, a trade-off between robustness of adult-survival to competition and reproduction gives coexistence. Invasion outcomes determined using (6) in (4a), using Supplemental (S20) and (S21) in (4b), and using numerical simulations in (4c). See Supplemental Figs. 3, 5 for verification of (4a), (4b) respectively using numerical simulations. Parameters are: (4a) *r*_*i*_ = *r*_*j*_ = 33.3, *m*_*j*_ = 18.5, *γ* = 2.5 and *α*_*j*_ = 7.6.; (4b) *r*_*j*_ = 5, *m*_*i*_ = *m*_*j*_ = 1, *γ* = 0.5, and *α*_*j*_ = 0.2.; (4c) *r*_*j*_ = 2.34, *m*_*i*_ = *m*_*j*_ = 0.4, *α*_*j*_ = 0.75 and *γ* = 1.

**Figure 5:**
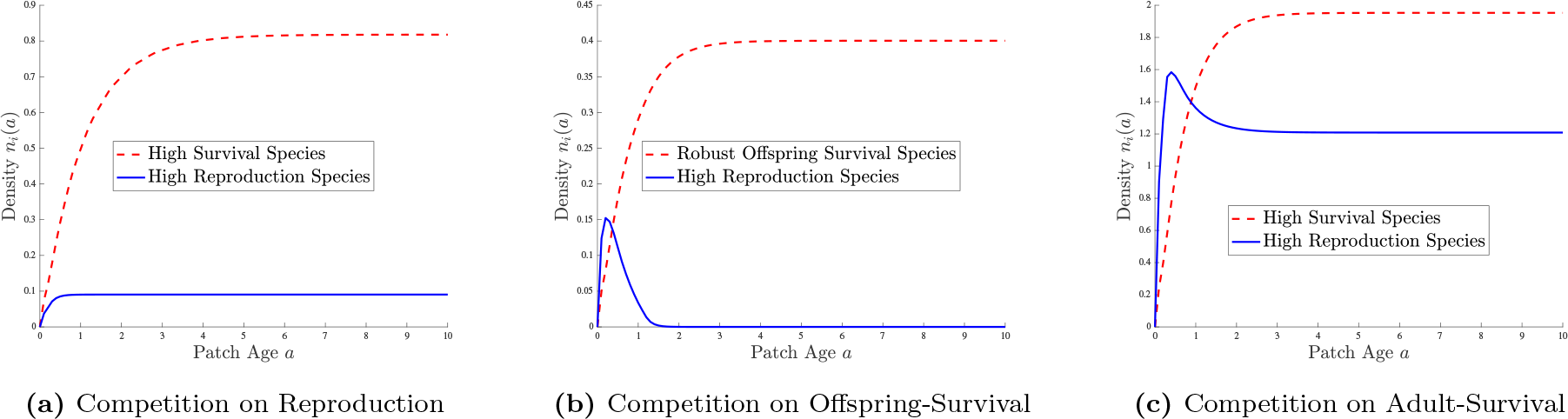
Coexistence May Not Involve “Successional Dynamics”. Dependence of species’ densities on patch-age at equilibrium when coexisting in each of our three model cases. In (5a) and (5c) the species are coexisting through a reproduction-adult-survival trade-off. In (5b) they are coexisting through a trade-off between reproduction and robustness of offspring-survival to competition. In (5a) and (5c) the high reproduction species remains at high density even in old patches. In (5a) it does not even lessen in abundance in old patches. In (5b) we see classical “successional dynamics”, with one species being more abundant in younger patches, but that species being excluded in older patches. All plots were generated using numerical simulations. Parameters are: (5a) *r*_*i*_ = 18, *m*_*i*_ = 7, *r*_*j*_ = 5.6, *m*_*j*_ = 1, *γ* = 1, *α*_*Rres*_ = 0, and *α*_*Rinv*_ = 0.; (5b) *r*_*i*_ = 35, *m*_*i*_ = *m*_*j*_ = 1, *r*_*j*_ = 5, *γ* = 1, *α*_*Rres*_ = 0.2, and *α*_*Rinv*_ = .325.; (5c) *r*_*i*_ = 10, *m*_*i*_ = *m*_*j*_ = 0.4, *r*_*j*_ = 2.34, *γ* = 1, *α*_*R*res_ = 0.75, and *α*_*R*inv_ = 8.

**Figure 6:**
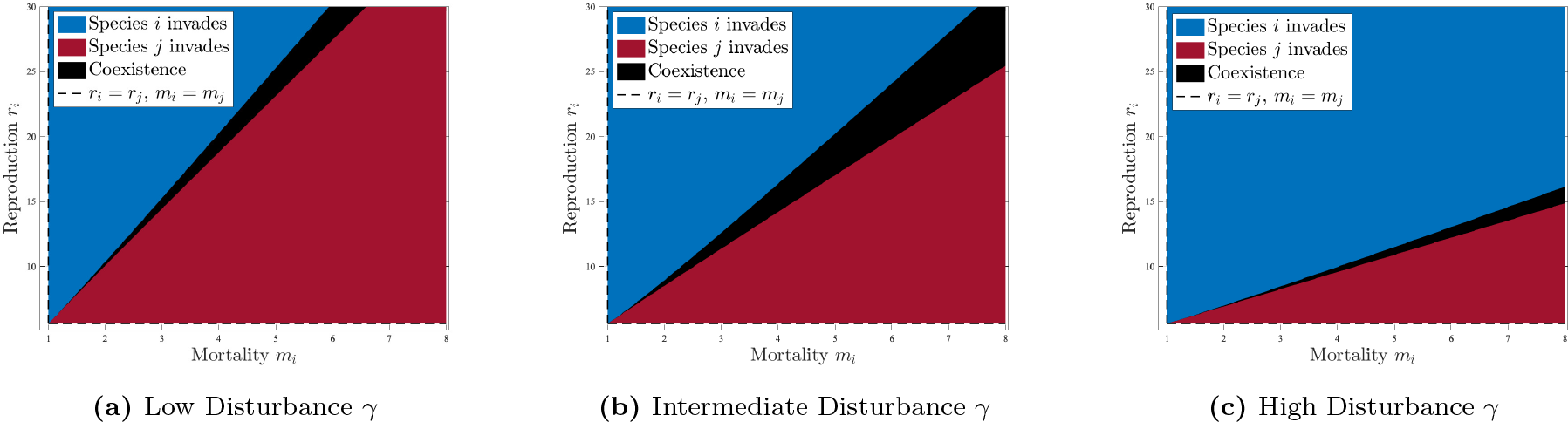
Intermediate Disturbance Yields Larger Coexistence Regimes. Shown are the coexistence regions for the competition acting on reproduction case, under a reproduction-survival trade-off, across three different disturbance rates. (See caption for Fig. 3 for more information about this type of coexistence figure.) We see that the range of parameters for species *i* (whose scale is constant across the plots) that also allow the two species to coexist, tends to be larger at intermediate values of the disturbance rate. This is just one example. See Supplementary Figs. 2, 4, 6, and 9 to see this trend for other model cases and trade-offs. Across all three plots, invasion outcomes were determined using (6), and *r*_*j*_ = 5.6, *m*_*j*_ = 1 and *α*_*i*_ = *α*_*j*_ = 1. Disturbance rate *γ* varies as (6a) *γ* = 0.1, (6b) *γ* = 0.4, (6c) *γ* = 2.5.

We note that a non-dimensionalized version of our model in the competition acting on reproduction case (see Supplementary Sec. 3.8) tells us that all model behavior, including coexistence, is determined only by the values of dimensionless parameters 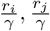 and 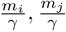 and the ratio 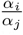. It is not determined by the disturbance rate *γ* in itself. This should also be true for our other model cases. However we chose not to plot results in this scaled parameter space because it clouds the dependence of the size of the coexistence regime on the disturbance rate *γ* (See Supplementary Sec. 3.8 for more details).

## Discussion

A key hypothesis of diversity maintenance is that disturbance opens up opportunities for species with different life history strategies to coexist. However, the development of general theory of life history trade-offs that can coexist under disturbance has been limited, not yet formulated in a way that captures the emergence of larger-scale competitive dynamics from within-patch population dynamics. Here we have presented a general partial differential equation model framework that accomplishes this, approaches to its analysis for key cases, and insights from carrying out those analyses, that together provide a starting point for further development of the theory of disturbance-generated coexistence.

In particular, this approach enables insights into how disturbance-generate coexistence arises from variation between species in their within-patch demographic strategy. A key prior model that provided general insights was focused on patch-level strategies (like the rate of colonization of new patches and whether a species out-competes others within a patch). Our framework instead considers species’ within-patch reproduction, offspring survival to adulthood, and adult survival, as well as the robustness of each of these components of within-patch demography to competition.

In addition to our work here providing a new model framework, we gained important insights from the analysis of key model cases. We discuss some of these and important areas of future work below.

### Within-Patch Demographic Trade-Offs versus Competition/Colonization

One insight is that variation between species on a simple demographic trade-off between reproduction and adult-survival can enable disturbance-generated coexistence. Even if the species experience the same proportional reduction in reproduction, or same proportional increase in mortality, due to competition, they can coexist—so long as one species tends to invest more in reproduction, and the other in adult-survival. We note that here “reproduction” is a per-capita rate of production of new *adults*—including the number of offspring produced—and their probability of survival to adulthood.

Life history theory takes as a given that organisms face a trade-off between reproduction and adult-survival, mediated by how their total investment in reproduction, or “reproductive effort” influences each [52, 53]. However, demonstrating this trade-off empirically can be challenging [54]. Though studies point to inter-specific variation on this trade-off (e.g. [55]), empirical information seems to be limited, with more information typically available regarding variation along a trade-off between offspring number and offspring-survival (e.g. [56]). Our work highlights the need for greater consideration of inter-specific variation along this trade-off in systems where disturbance-generated coexistence is the focus. Indeed, recent work shows previously unexplored variation among tropical tree species from being long-lived to having high recruitment rates [57] that appears analogous to the variation on this trade-off in our model.

We note some similarity—but important differences—between coexistence through this reproduction versus adult-survival trade-off and the classical “competition-colonization” trade-off [14, 17–19]. The similarity is that the better reproducing species in our model would arguably be a better colonizer of new patches, and the better surviving species in our model is in fact the better competitor for a patch if it were isolated (See Supplemental Sec. 2). However, in our model, the better reproducer and the better competitor both arrive to patches in small density immediately. In contrast, the competition-colonization trade-off model implicitly incorporates a possible delay in patches being occupied by either species, through a focus on the proportion of patches being occupied versus empty, or only be occupied by the better colonizer.

Also, in our model when there is coexistence through a reproduction versus adult-survival trade-off, both species are present at high abundances in older patches. In other words, the coexistence does not involve classical “successional dynamics” (where the better competitor species immediately captures patches from the better colonizer) as presumed in the competition-colonization trade-off model. Thus there is no clear mapping between coexistence through this trade-off in our model and the competition-colonization trade-off.

However, one of the trade-offs with “competition-robustness” that we found to be coexistence-generating may have a clearer mapping with the competition-colonization trade-off, specifically the trade-off between reproduction and robustness-of-offspring-survival to competition. When there is coexistence through variation along this trade-off in our model, the more robust competitor indeed out-competes the better reproducer in older patches, even patches remaining part of the landscape, because it blocks the better reproducer from recruiting in older patches. So this trade-off involves dynamics as a function of patch-age that are consistent with the competition-colonization trade-off, and might be viewed as a more specific example of it.

Another important note is that while the competition-colonization trade-off has been to some degree dismissed as a potential mechanism maintaining seed-size diversity [58] (which can be substantial among coexisting species [59]), our reproduction versus robustness-to-competition trade-off provides a more specific trade-off that is still plausible. The competition-colonization trade-off was dismissed based on empirical studies suggesting “competition is not particularly asymmetric in favor of large-seeded species” (which would be the species with lower fecundity or colonization rates). However, those empirical studies were focused on competition among seedlings [60, 61], not competitive effects from adults, the key aspect of competition in which the species vary in our trade-off. There is some association between seed-size and shade-tolerance among tree species, and hence evidence for variation among species on a reproduction versus offspring-survival competition robustness [60]. Indeed, heterogeneity in availability of light and nutrients, created by competition with established adults, is presented as an example mechanism creating the variation in stress on the landscape required for the alternative “tolerance-fecundity” trade-off explanation of seed-size diversity [58]. Our trade-off can also be viewed as similar to that alternative trade-off, but where stress variation is disturbance-generated and dynamic rather than having a fixed distribution on the landscape. However, stress to seedlings will have other sources of variation in forests as well, validating consideration of the trade-off with tolerance in a broad way.

A final note related to the competition-colonization trade-off is that an important contrast exists between the insights provided in our model about what variation in demographic strategy enables coexistence, and insights that result from interpreting the competition-colonization trade-off model as describing occupancy of “patches” that are each the scale of a single individual [22]. This alternative interpretation of the competition-colonization trade-off model takes the presumption that the offspring of a better competitor can take over a site occupied by an adult of a poorer competitor. Although this may happen in the long term in the patch for some organisms, the model does not provide more detail of what individual-scale demographic strategy variation constitutes the better competitor strategy, so it is difficult to relate to our framework. Perhaps a more detailed version of that model, keeping track of the dynamics of replacement events at the individual scale, rather than assuming the occur immediately, could clarify. Also worth noting is that when viewing competition-colonization trade-off model as describing the scale of individuals, one can think of it as just a particular way of writing Lotka-Volterra competitive dynamics—one with a competitive hierarchy. From that perspective, spatial structure is not essential to the stable coexistence that it produces. Instead what is key is a trade-off between how much more sensitive a species is to competition from the other (compared to competition from itself) and either fecundity or morality, as pointed out previously [62]. The coexistence we consider here differs from that, as it indeed requires disturbance and patch structure.

### Easier Coexistence at Intermediate Disturbance

The original proposal that disturbance can generate coexistence of competing species [10] has been at times termed the “Intermediate Disturbance Hypothesis” (IDH), to emphasize the related prediction that the diversity maintained by this mechanism would be highest when disturbance is not too frequent nor too infrequent. Recently there has been discussion regarding the impact of IDH as a coexistence mechanism [63]. However, this was a critique of the misconception that disturbance can in itself generate stable coexistence of species by reducing species densities, rather than the idea that disturbance can open up the opportunity for species differing in life history to coexist, which is our focus here. We found in our model cases that coexistence regimes do tend to be larger at intermediate disturbance rates. Further study of our model in a multi-species formulation is needed to determine if this translates into the stable coexistence of a larger number of types being possible at intermediate disturbance rates. However, empirical examination of the IDH has considered disturbance that is “intermediate” in a variety of senses, including intermediate times since a patch has been disturbed and intermediate intensity of disturbance as measured by how much biomass is removed, a complexity supported by simulation studies [64]. Likely a similar complexity would arise in our model, but it is clear from our analyses that disturbance rate is indeed an important factor of the ease with which species differing in demographic strategy can coexist.

### Coexisting Life History Variation May Vary Across Systems

We also found that the specific coexistence opportunities that become available under disturbance depend on what aspects of demography are most impacted by competition. In particular, a reproduction versus adult-survival trade-off did not produce coexistence when competition acts on offspring-survival alone. The action of competition-on-reproduction and adult-survival is needed for it to be coexistence-generating. Furthermore, coexistence through trade-offs with robustness to competition of the various components of demography require the presence of an influence of competition on that aspect of demography. Which aspects of demography are most strongly impacted by competition likely varies across systems. Our results could provide insight into variation across those systems in the types of life history strategies that coexist.

Negative density-dependence of reproduction—at least in forests—is difficult to measure, as it would require measuring the total seed production of specific adult trees, which is challenging. Our work highlights the need for measurements of this competition in order to build accurate models of disturbance-generated tree species coexistence. Adult-survival may not be strongly affected by competition for tree species in forests. Mortality of tropical trees has been found to be relatively independent of diameter among larger individuals (a proxy for adulthood) [65]. Size is a rough proxy of the amount of competition an individual may be experiencing (with smaller trees experiencing more competition through shading). In contrast, competitive pressure from established adults acts strongly on offspring-survival for trees, since seedlings and saplings are inhibited when in the understory of large adults.

### Important Areas of Future Work

There are a great number of ways in which this work could be extended. Especially pressing are the need to: a) carry out analyses for multiple species, b) consider—in concert—the action of competition on multiple components of demography, c) consider variation between species in the time it takes to become adults (or alternatively in the”age of first reproduction”), and d) consider variation in distance between patches and variation in dispersal distance between species. These may all be challenging to carry out analytically. Factoring in an “age of first reproduction” would require considering time delays between when reproduction occurs and when it contributes to the adult population in patches. As is, we only obtained the two species structured equilibrium analytically for one model case, and that equilibrium is needed to consider at least the invasion conditions for a third species. Also, competition acting on adult-survival is difficult to make any analytical progress with. However, recently, empirically supported approximations for forests have been used to make analytical progress with forest-specific models incorporating size-structure [28, 66] and even a combination of patch-age and size-structure [32]. This could perhaps point to useful limits in the exploration of the general model framework we present. Such an exploration could also reveal how these important forest-focused results might be understood in terms of our framework, a question also worth considering for other forest-specific and more general disturbance and succession models that have been studied through simulation [27, 67, 68]. Careful exploration of our model in parameter space using numerical simulations may also be fruitful for identifying parameter combinations leading to coexistence in more complex scenarios. An ultimate goal is to understand how demographic parameters combine to determine species’ separation in the “niche” space [69] created by disturbance. For this it may be useful to draw on approaches to quantifying the robustness of coexistence, which will be inversely related to these “niche differences” in terms of species differences when interacting with regulating factors [7, 70].

One final note is that our model framework is not currently designed to incorporate stochasticity, which can have profound effects on community structure [71]. Indeed, recent work has shown that the persistence of species with complementary life history strategies is enhanced in a stochastic context, without the presence of disturbance that would enable their stable coexistence [72]. Other works have shown that patterns of variation in abundance with life history strategies can arise in that context as well [73, 74]. Stochastic effects could also open up the opportunity for mechanisms of stable coexistence, and would be an important extension of our framework in future.

### Conclusions

In conclusion, our work here provides a new tool for further development of the theory of disturbance-generated coexistence, a tool which allows for theory based on within-patch demographic strategies. The development of general theory using this tool will reveal new more detailed mechanisms of disturbance-generated coexistence, as well as deeper insight into known mechanisms, and serves as a complement to system-focused approaches, providing guidance on what model ingredients are essential, and ideally, some understanding of the patterns of coexistence across systems where disturbance plays a key role.

## Source Code

Source code for simulations and analytical work is available at https://github.com/utrigos/Disturbance_Generated_Competitive_Coexistence_Code.

## 1 Acknowledgements

The research of MGD was partially supported by NSF-DMS-2205937 and NSF-DMS RTG 18403. RL acknowledges the financial support of Carl Tryggers Stieftelse via the grant CTS 21:1656. The research of AO was supported by NSF-DEB-2122309. We thank Charlie Doering for his insightful suggestions on this research.

