## Supplemental Information for "Disturbance-generated competitive coexistence"

### Supplemental information for
#### Disturbance-generated competitive coexistence

Ursula A. Trigos-Raczkowski, Rainey Lyons, Matias G. Delgadino, Azmy S. Ackleh, Annette Ostling

November 2, 2023

### Contents

**1 Dynamics of $\rho$ 1**

**2 Coexistence Outcomes in the Absence of Disturbance and Succession 2**
2.1 Competition Acting on Reproduction and Offspring-Survival . . . . . 2
2.2 Density-Dependent Adult-Survival Case . . . . . 3

**3 Competition Acting on Reproduction 3**

**5 Competition Acting on Adult-Survival 18**

**6 Numerical Method 18**

### 1 Dynamics of $\rho$

Our patch dynamics are described by

$$\frac{\partial\rho(a,t)}{\partial t} = -\frac{\partial\rho(a,t)}{\partial a} - \gamma(a)\rho(a,t)$$
$$\rho(0,t) = \int_0^\infty \gamma(a)\rho(a,t)da.$$

For all of our model cases we take the simplifying assumption that the disturbance rate is constant with respect to patch age, i.e. $\rho(a) = \rho$, in which case the steady state for our patch dynamics is

$$\tilde{\rho}(a) = \gamma e^{-\gamma a}. \tag{S1}$$

Though the full dynamics of this McKendrick–Von Foerster style model are well known, we demonstrate the local stability of this equilibrium below for the reader's benefit.

We examine the dynamics of $\rho$ which satisfy equation (MF1)-(MF2). In particular, we are interested under which conditions $\rho$ converges to the steady-state, $\rho^*(a)$ given by equation (S1). We assume $\gamma(a) \equiv \gamma > 0$. We first begin by noting $P(t) := \int_0^\infty \rho(a,t)$ is constant in time. Indeed, integrating (MF1) over $a$ yields

$$\frac{d}{dt}P(t) = -(\rho(a,t)|_0^\infty) - \int_0^\infty \gamma(a)\rho(a,t)da = 0.$$

From this, it is clear that if the initial condition, $\rho_0(a)$, is such that $\int_0^\infty \rho_0(a)da \neq 1$, $\rho$ will not converge to the steady-state.

Let us assume now that $\int_0^\infty \rho_0(a)da = 1$. Then, the function $\omega(a,t) := \rho(a,t) - \tilde{\rho}(a)$ satisfies the following equation:

$$\begin{cases} \partial_t \omega(a,t) + \partial_a \omega(a,t) = -\gamma\omega \\ \omega(0,t) = \int_0^\infty \gamma\rho(a,t)da - \gamma = 0 \\ \omega(a,0) = \rho_0(a) - \tilde{\rho}(a) =: \omega_0(a) \end{cases} . \tag{S2}$$

Following standard characteristic arguments, we conclude the solution of equation (S2) is given by

$$\omega(a,t) = \begin{cases} \omega_0(a-t)\exp(-\gamma t) & a \geq t \\ 0 & t > a \end{cases} .$$

Which uniformly goes to 0 as $t \longrightarrow \infty$.

### 2 Coexistence Outcomes in the Absence of Disturbance and Succession

The addition of disturbance and patch-age structure in the model we present are crucial in the coexistence of the species. Here we show that without disturbance and the patch-age structure it creates, coexistence is simply not possible in our model unless we allow for differences in the competitive effects each species exercises on the other.

Removing disturbance and the patch-age structure it creates is equivalent to narrowing our landscape to a single patch. We derive competitive outcomes for this simplified version of our model cases below.

#### 2.1 Competition Acting on Reproduction and Offspring-Survival

In both the competition acting on reproduction and competition acting on offspring-survival cases of our model, removing disturbance and thus patch-age structure from the population dynamic equations would yield (in this case the patch-level densities $ni(a)$ and $n_j(a)$ are independent of $a$ so we denote them simply by $n_i$ and $n_j$):

$$\begin{aligned} \frac{dn_i}{dt} &= r_i n_i(1 - \alpha_i n_i - \alpha_i n_j) - m_i n_i \\ \frac{dn_j}{dt} &= r_j n_j(1 - \alpha_j n_i - \alpha_j n_j) - m_j n_j. \end{aligned}$$

Rearranging these to be in the form in which Lotka-Volterra competition equations are typically written we obtain:

$$\begin{aligned} \frac{dn_i}{dt} &= (r_i - m_i)n_i\left(1 - \frac{r_i\alpha_i}{r_i - m_i}n_i(t) - \frac{r_i\alpha_i}{r_i - m_i}n_j(t)\right) \\ \frac{dn_j}{dt} &= (r_j - m_j)n_j\left(1 - \frac{r_j\alpha_j}{r_j - m_j}n_j(t) - \frac{r_j\alpha_j}{r_j - m_j}n_j(t)\right). \end{aligned}$$

These can be used to derive the conditions for species $i$ to invade species $j$ ($\frac{1}{n_i}\frac{dn_i}{dt}|_{n_i \approx 0, n_j *} > 0$) and the condition for species $j$ to invade species $i$ ($\frac{1}{n_j}\frac{dn_j}{dt}|_{n_i *, n_j \approx 0} > 0$), resulting in the conditions below respectively:

$$\frac{r_i\alpha_i}{r_i - m_i} < \frac{r_j\alpha_j}{r_j - m_j} \quad \text{and} \quad \frac{r_j\alpha_j}{r_j - m_j} < \frac{r_i\alpha_i}{r_i - m_i}.$$

These conditions are the reverse of each other and clearly cannot hold simultaneously. Thus coexistence in the sense of mutual invasion is simply impossible without the addition of disturbance and patch dynamics (unless we vary the competitive effects).

Note that if instead we considered a case of the model where the competitive effects on species $i$ depend on whether it is affected by species $i$ or $j$ that model for a single patch would look like

$$\begin{aligned} \frac{dn_i}{dt} &= r_i n_i(1 - \alpha_{ii} n_i - \alpha_{ij} n_j) - m_i n_i \\ \frac{dn_j}{dt} &= r_j n_j(1 - \alpha_{ji} n_i - \alpha_{jj} n_j) - m_j n_j. \end{aligned}$$

In that case, in order to have coexistence (in the sense of mutual invasion) we would require

$$\frac{r_i\alpha_{ij}}{r_i - m_i} < \frac{r_j\alpha_{jj}}{r_j - m_j} \quad \text{and} \quad \frac{r_j\alpha_{ji}}{r_j - m_j} < \frac{r_i\alpha_{ii}}{r_i - m_i}.$$

Even if species are identical in their reproduction and mortality parameters, coexistence in this system can occur iff $\alpha_{ij} < \alpha_{jj}$ and $\alpha_{ji} < \alpha_{ii}$, i.e. when interspecific competition is less than intraspecific competition, and thus coexistence can occur without the disturbance or patch-age dynamics in this special case. Our framework could eventually be used to study how disturbance and patch-dynamics modify such coexistence generated by classical *within-patch* "niche differences" (i.e. differential use of within-patch resources).

#### 2.2 Density-Dependent Adult-Survival Case

In the competition acting on adult-survival case, removing disturbance and patch dynamics from our model equations would result in

$$\frac{dn_i}{dt} = r_i n_i - m_i n_i (1 - \alpha_i n_i - \alpha_i n_j)$$
$$\frac{dn_j}{dt} = r_j n_j - m_j n_j (1 - \alpha_j n_i - \alpha_j n_j).$$

Rearranging these to be in the form in which Lotka-Volterra competition equations are typically written we obtain:

$$\frac{dn_i}{dt} = (r_i - m_i)\, n_i \left(1 + \frac{m_i \alpha_i}{(r_i - m_i)} n_i + \frac{m_i \alpha_i}{(r_i - m_i)} n_j\right)$$
$$\frac{dn_j}{dt} = (r_j - m_j)\, n_j \left(1 + \frac{m_j \alpha_j}{(r_j - m_j)} n_i + \frac{m_j \alpha_j}{(r_j - m_j)} n_j\right).$$

The conditions for species $i$ to invade species $j$ ($\frac{1}{n_i}\frac{dn_i}{dt}|_{n_i \approx 0, n_j*} > 0$) and the condition for species $j$ to invade species $i$ ($\frac{1}{n_j}\frac{dn_j}{dt}|_{n_i*, n_j \approx 0} > 0$) are then respectively:

$$\frac{m_i \alpha_i}{r_i - m_i} < \frac{m_j \alpha_j}{r_j - m_j} \quad \text{and} \quad \frac{m_j \alpha_j}{r_j - m_j} < \frac{m_i \alpha_i}{r_i - m_i},$$

and we can clearly see that these cannot be satisfied simultaneously. Hence coexistence in the sense of mutual invasion cannot occur without disturbance and patch dynamics.

If we instead consider a case without patch-dynamics but allowing for differences in competitive effects we would have

$$\frac{dn_i}{dt} = r_i n_i - m_i n_i (1 - \alpha_{ii} n_i - \alpha_{ij} n_j)$$
$$\frac{dn_j}{dt} = r_j n_j - m_j n_j (1 - \alpha_{ji} n_i - \alpha_{jj} n_j).$$

In this case mutual invasion would require

$$\frac{m_i \alpha_{ii}}{r_i - m_i} < \frac{m_j \alpha_{ji}}{r_j - m_j} \quad \text{and} \quad \frac{m_j \alpha_{jj}}{r_j - m_j} < \frac{m_i \alpha_{ij}}{r_i - m_i}.$$

### 3 Competition Acting on Reproduction

The model equations for this case are

$$\frac{\partial n_i(a,t)}{\partial t} = \underbrace{r_i \Big( \int_0^\infty \rho(a',t)\, n_i(a',t) \max[0, (1 - \alpha_i n_i(a,t) - \alpha_i n_j(a,t))]\, da' \Big)}_{\text{recruitment}} - \underbrace{m_i n_i(a,t)}_{\text{mortality}} - \underbrace{\frac{\partial n_i(a,t)}{\partial a}}_{\text{patch-aging}}$$

$$\frac{\partial \rho(a,t)}{\partial t} = -\frac{\partial \rho(a,t)}{\partial a} - \gamma \rho(a,t)$$
$$\rho(0,t) = \int_0^\infty \gamma \rho(a,t) da$$
$$n_i(0,t) = 0.$$

Note that the integral in the recruitment term represents the summation of reproduction across all patches of all ages, modified by negative density-dependence in each patch, and $r_i$ reflects a combination of the baseline rate of reproduction in an empty patch, as well as the rate of offspring survival in the patches where they land (assumed to be density-independent in this model).

The following expression for the equilibrium density of species $i$ can be derived in either the case where it is on its own or in the presence of species $j$:

$$\tilde{n}_i(a) = \frac{r_i \tilde{S}_i}{m_i} \left(1 - e^{-m_i a}\right), \tag{S3}$$

where $\tilde{S}_i$ is the integral in the recruitment term at equilibrium. We denote the density of species $i$ on its own at equilibrium as $\tilde{n}_{i|0}$ and in the presence of species $j$ as $\tilde{n}_{i|j}$ (and for those cases the appropriate reproduction integral by $\tilde{S}_{i|0}$ and $\tilde{S}_{i|j}$). For example, as stated in the main text, we have

$$\tilde{S}_{i|j} = \int_0^\infty \rho(a') \tilde{n}_{i|j}(a') \max[0, \big(1 - \alpha_i \tilde{n}_{i|j}(a') - \alpha_i \tilde{n}_{j|i}(a')\big)] da'.$$

#### 3.1 Obtaining Analytical Results and the Max Function

Note that (S3) is a monotonically increasing function of $a$, with $n_i(0) = 0$ (as required by our boundary condition). Due to this, we know that at equilibrium the max function in the recruitment term will yield 1 at $a = 0$ and decline from there, possibly giving 0 at large $a$. Hence the max function will at the most, impose an upper limit on the integral. We carry out analytical calculations for this case below presuming we are in a parameter regime where the max function imposes no limitations on the integral and hence it can be taken to infinity. We were not able to solve analytically for an expression for critical patch-age and hence were also unable to determine analytical conditions for coexistence. However, when carrying out numerical simulations to verify our analytical results, we also check that any analytical coexistence-generating trade-offs do indeed overlap in parameter space with regimes where species densities remain small enough at large patch-ages that our analytical approach is indeed valid, (i.e. are small enough that we can disregard the max function because the values would not drive reproduction to zero or negative values).

#### 3.2 One Species Equilibrium

At equilibrium we solve for the $\tilde{S}_{i|0}$ value using

$$\tilde{S}_{i|0} = \int_0^\infty \gamma e^{-\gamma a'} \frac{r_i \tilde{S}_{i|0}}{m_i}\left(1 - e^{-m_i a'}\right)\left(1 - \alpha_i \frac{r_i \tilde{S}_{i|0}}{m_i}\left(1 - e^{-m_i a'}\right)\right) da',$$

resulting in

$$\tilde{S}_{i|0} = \frac{(\gamma + 2m_i)(r_i - \gamma - m_i)}{2\alpha_i r_i^2},$$

and hence the density of species $i$ at equilibrium on its own follows:

$$\tilde{n_i}(a) = \frac{(\gamma + 2m_i)(r_i - \gamma - m_i)}{2\alpha_i r_i m_i}\left(1 - e^{-m_i a}\right).$$

The expressions for species $j$ are analogous.

#### 3.3 Two Species Coexistence Equilibrium and its Feasibility

We derive the two species coexistence equilibrium constants for species $i$ and $j$, ($\tilde{S}_{i|j}$, and $\tilde{S}_{j|i}$) by solving the coupled equations:

$$\tilde{S}_{i|j} = \int_0^\infty \gamma e^{-\gamma a'} \frac{r_i \tilde{S}_{i|j}}{m_i}\left(1 - e^{-m_i a'}\right)\left(1 - \alpha_i \frac{r_i \tilde{S}_{i|j}}{m_i}\left(1 - e^{-m_i a'}\right) - \alpha_i \frac{r_j \tilde{S}_{j|i}}{m_j}\left(1 - e^{-m_j a'}\right)\right) da'$$

$$\tilde{S}_{j|i} = \int_0^\infty \gamma e^{-\gamma a'} \frac{r_j \tilde{S}_{j|i}}{m_j}\left(1 - e^{-m_j a'}\right)\left(1 - \alpha_j \frac{r_i \tilde{S}_{i|j}}{m_i}\left(1 - e^{-m_i a'}\right) - \alpha_j \frac{r_j \tilde{S}_{j|i}}{m_j}\left(1 - e^{-m_j a'}\right)\right) da'.$$

Focusing on their non-zero solutions, we drive the similar expressions

$$\tilde{S}_{i|j} = \frac{(\gamma + m_i)(\gamma + 2m_i)(\gamma + m_j)(\gamma + 2m_j)(\gamma + m_i + m_j)^2}{\gamma r_i r_j \alpha_i \alpha_j (m_i - m_j)^2 (3\gamma + 2m_i + 2m_j)}\left(\frac{2r_j\alpha_j(r_i - \gamma - m_i)}{r_i(\gamma + 2m_j)} - \frac{\alpha_i(r_j - \gamma - m_j)(2\gamma + m_i + m_j)}{(\gamma + m_j)(\gamma + m_i + m_j)}\right)$$

$$\tilde{S}_{j|i} = \frac{(\gamma + m_j)(\gamma + 2m_j)(\gamma + m_i)(\gamma + 2m_i)(\gamma + m_j + m_i)^2}{\gamma r_j r_i \alpha_j \alpha_i (m_j - m_i)^2 (3\gamma + 2m_j + 2m_i)}\left(\frac{2r_i\alpha_i(r_j - \gamma - m_j)}{r_j(\gamma + 2m_i)} - \frac{\alpha_j(r_i - \gamma - m_i)(2\gamma + m_i + m_j)}{(\gamma + m_i)(\gamma + m_i + m_j)}\right).$$

For species $i$ and $j$ to both have feasible populations, we need $\tilde{S}_{i|j}$, and $\tilde{S}_{j|i}$ to be positive. We can see the coefficients out front are always positive so long as the species differ in their mortality rates, (i.e. $m_i \neq m_j$), so feasible coexistence requires the species to differ in their mortality rates. Additionally we need the terms in parenthesis to be positive.

Hence for species $i$ to have a feasible population at the coexistence equilibrium, i.e. for $\tilde{S}_{i|j} > 0$ we find the following condition:

$$\frac{\frac{r_i\alpha_i}{(r_i - (\gamma + m_i))}}{\frac{r_j\alpha_j}{(r_j - (\gamma + m_j))}} < \frac{2(\gamma + m_j)(\gamma + m_i + m_j)}{(2\gamma + m_i + m_j)(\gamma + 2m_j)}. \tag{S4}$$

For species $j$ to have a feasible population at the coexistence equilibrium, i.e. for $\tilde{S}_{j|i} > 0$ we find the comparable condition:

$$\frac{(\gamma + 2m_i)(2\gamma + m_i + m_j)}{2(\gamma + m_i)(\gamma + m_i + m_j)} < \frac{\frac{r_i\alpha_i}{(r_i - (\gamma + m_i))}}{\frac{r_j\alpha_j}{(r_j - (\gamma + m_j))}}. \tag{S5}$$

Finally, the conditions for a feasible coexistence equilibrium can be summarized as

$$\frac{(\gamma + 2m_i)(2\gamma + m_i + m_j)}{2(\gamma + m_i)(\gamma + m_i + m_j)} < \frac{\frac{r_i\alpha_i}{r_i - (\gamma + m_i)}}{\frac{r_j\alpha_j}{r_j - (\gamma + m_j)}} < \frac{2(\gamma + m_j)(\gamma + m_i + m_j)}{(2\gamma + m_i + m_j)(\gamma + 2m_j)}.$$

#### 3.4 Mutual Invasion Analysis

Here we provide the details of the invasion analysis for the competition acting on reproduction model. First we derive dynamical equations for the average density of species $i$ across patches of all ages. We begin with our PDE for $n_i(a)$:

$$\frac{\partial n_i(a,t)}{\partial t} = r_i \int_0^\infty \rho(a',t)\, n_i(a',t) f_i\big(n_i(a',t), n_j(a',t)\big) da' - m_i n_i(a,t) - \frac{\partial n_i(a,t)}{\partial a},$$

where for efficiency we wrote the density-dependence on the reproduction term as following a general functional form $f_i\big(n_i(a',t), n_j(a',t)\big)$. Now we define $g_i(a,t) \equiv n_i(a,t)\ \rho(a,t)$ and derive its dynamical equation, which we can then integrate over to get a dynamical equation for the average density across patches of species $i$, $N_i(t)$, defined as

$$N_i(t) = \int_0^\infty n_i(a,t)\rho(a,t)da.$$

We obtain this dynamical equation using the product rule for derivatives and substituting the dynamical equation for $\rho(a,t)$.

$$\begin{aligned}\frac{\partial g_i(a,t)}{\partial t} &= \left(r_i \int_0^\infty \rho(a',t)\ n_i(a',t) f_i\big(n_i(a',t), n_j(a',t)\big)da' - m_i n_i(a,t) - \frac{\partial n_i(a,t)}{\partial a}\right)\rho(a,t) + n_i(a,t)\left(-\frac{\partial \rho(a,t)}{\partial a} - \gamma\rho(a,t)\right)\\ &= r_i\Big[\int_0^\infty \rho(a',t)\ n_i(a',t) f_i\big(n_i(a',t), n_j(a',t)\big)da'\Big]\rho(a,t) - m_i\ g_i(a,t) - \frac{\partial g_i(a,t)}{\partial a} - \gamma\ g_i(a,t)\end{aligned}$$

Integrating over all patches we obtain the following dynamical equation for $N_i(t)$:

$$\frac{dN_i}{dt} = \int_0^\infty \rho(a,t)da\ \left(r_i \int_0^\infty \rho(a',t)\ n_i(a',t) f_i\big(n_i(a',t), n_j(a',t)\big)da'\right) - (m_i + \gamma)\, N_i(t) - \left(\lim_{a\to\infty} g_i(a,t) - g_i(0,t)\right).$$

We then assume $g_i(a,t)$ is zero at $a = 0$ (because our boundary condition is $n_i(0,t) = 0$ and $\rho(0,t)$ is finite), that $\lim_{a\to\infty} n_i(a,t) = 0$ (because $\lim_{a\to\infty} \rho(a,t) = 0$ will hold once the patch dynamics come to equilibrium) as well as the fact that $n_i(a)$ is finite at large patch-ages. We then divide both sides by the total density to give us an expression for the per-capita total density time derivative of species $i$ that is useful for invasion analysis.

$$\frac{1}{N_i}\frac{dN_i}{dt} = \frac{1}{N_i}\left(r_i \int_0^\infty \rho(a',t)\ n_i(a',t) f_i\big(n_i(a',t), n_j(a',t)da'\big)\right) - (m_i + \gamma).$$

##### 3.4.1 Invader Density Constant Across Patch-Age

We also derived the invasion growth rate assuming a constant distribution across patch-ages of the invader. This would correspond to an initial small density of individuals of the invading species spread uniformly across patches of different ages.

$$\frac{1}{N_i}\frac{dN_i}{dt}\Big|_{\epsilon_i(a)\approx\epsilon, \tilde{n}_{j|0}(a)} = r_i \frac{\int_0^\infty da\ \rho(a,t)\ \not{\epsilon}\ f_i\big(\epsilon, \tilde{n}_{j|0}(a)\big)}{\underbrace{\int_0^\infty da\ \not{\epsilon}\rho(a)}_{1}} - (m_i + \gamma) > 0.$$

This results in

$$\frac{1}{N_i}\frac{dN_i}{dt}\Big|_{\epsilon_i(a)\approx\epsilon, \tilde{n}_{j|0}(a)} = r_i \int_0^\infty da\ \rho(a)\left(1 - \epsilon - \tilde{n}_{j|0}\right) - (m_i + \gamma) > 0.$$

When $\epsilon << 1$:

$$\begin{aligned}\frac{1}{N_i}\frac{dN_i}{dt}\Big|_{\epsilon_i(a)\approx\epsilon, \tilde{n}_{j|0}(a)} &\approx r_i \int_0^\infty da\ \rho(a)\left(1 - \tilde{n}_{j|0}\right) - (m_i + \gamma) > 0\\ &= r_i\left(1 - \tilde{N}_{j|0}\right) - (m_i + \gamma) > 0,\end{aligned}$$

where $\tilde{N}_{j|0}$ is the equilibrium landscape average density of species $j$ when on its own. We can obtain this using the equilibrium density for species $j$ on its own, i.e.:

$$\tilde{n}_{j|0}(a) = \frac{r_j \tilde{S}_{j|0}}{m_j}\left(1 - e^{-m_j a}\right).$$

Resulting in

$$\tilde{N}_{j|0} = \int_0^\infty da\ \rho(a)\tilde{n}_{j|0}(a) = \frac{r_j\gamma\tilde{S}_{j|0}}{m_j}\left(\frac{m_j}{\gamma\left(\gamma + m_j\right)}\right).$$

We know that our value of $\tilde{S}_{j|0}$ (see Sec. 3.2 for details) is

$$\tilde{S}_{j|0} = \frac{\left(\gamma + 2m_j\right)\left(r_j - \gamma - m_j\right)}{2r_j^2}.$$

Substituting, we obtain

$$\tilde{N}_{j|0} = \frac{\left(r_j - \gamma - m_j\right)\left(\gamma + 2m_j\right)}{2r_j\left(\gamma + m_j\right)}.$$

The invasion condition is then

$$\frac{2r_j\left(\gamma + m_j\right) - \left(r_j - \gamma - m_j\right)\left(\gamma + 2m_j\right)}{2r_j\left(\gamma + m_j\right)} > \frac{m_i + \gamma}{r_i}. \tag{S6}$$

The condition for $j$ to invade $i$ can be written analogously.

$$\frac{2r_i\left(\gamma + m_i\right) - \left(r_i - \gamma - m_i\right)\left(\gamma + 2m_i\right)}{2r_i\left(\gamma + m_i\right)} > \frac{m_j + \gamma}{r_j}. \tag{S7}$$

These conditions end up giving a larger region of mutual invasion than the region of mutual invasion when we assume that the invader abundance over patch-ages is proportional to its equilibrium form. As described in the main text, we found with numerical simulations that indeed in this wider parameter regime the invader would initially increase in its landscape abundance if started with a constant abundance over patch-ages. However, its structure over patch-ages would quickly go to its equilibrium form, and then the species would decline to extinction unless parameters are in the narrower regime where coexistence is feasible, and invasion would be predicted if the invader density is assumed proportional to this form.

##### 3.4.2 Invader Density Across Patch-Ages Assumed to Quickly Go To its Equilibrium Form

Here we derive an analytical expression for the invasion growth rate in the case where we assume the invader's initial density quickly becomes approximately equal to the form it would at equilibrium, i.e. assuming $\epsilon_i(a) \approx \epsilon\,(1 - e^{-m_i a})$.

$$\frac{1}{N_i}\frac{dN_i}{dt}\Big|_{\epsilon_i(a),\tilde{n}_{j|0}(a)} = r_i \frac{\int_0^\infty \rho(a,t)\ \epsilon_i(a)\ f_i\big(\epsilon_i(a), \tilde{n}_{j|0}(a)\big)da}{\int_0^\infty \epsilon_i(a)\ \rho(a)da} - (m_i + \gamma) > 0 \tag{S8}$$

$$\frac{1}{N_i}\frac{dN_i}{dt}\Big|_{\epsilon_i(a),\tilde{n}_{j|0}(a)} \approx r_i \frac{\int_0^\infty \rho(a,t)\,\cancel{\epsilon}\ (1 - e^{-m_i a})\ f_i\big(\epsilon_i(a), \tilde{n}_{j|0}(a)\big)da}{\int_0^\infty \cancel{\epsilon}\ (1 - e^{-m_i a})\ \rho(a)da} - (m_i + \gamma) > 0$$

$$\frac{1}{N_i}\frac{dN_i}{dt}\Big|_{\epsilon_i(a),\tilde{n}_{j|0}(a)} \approx \frac{r_i \int_0^\infty e^{-\gamma a}\ \ (1 - e^{-m_i a})\ f_i\big(\epsilon_i(a), \tilde{n}_{j|0}(a)\big)da}{\int_0^\infty e^{-\gamma a}\ \ (1 - e^{-m_i a})\,da} - (m_i + \gamma) > 0.$$

For linear $f_i$, , we have

$$\frac{1}{N_i}\frac{dN_i}{dt}\Big|_{\epsilon_i(a),\tilde{n}_{j|0}(a)} \approx \frac{r_i \int_0^\infty e^{-\gamma a}\ \ (1 - e^{-m_i a})\ \big(1 - \alpha_i\epsilon_i(a) - \alpha_i\tilde{n}_{j|0}(a)\big)da}{\frac{m_i}{\gamma(\gamma+m_i)}} - (m_i + \gamma) > 0.$$

For $\epsilon << 1$

$$\frac{1}{N_i}\frac{dN_i}{dt}\Big|_{\epsilon_i(a),\tilde{n}_{j|0}(a)} \approx \frac{r_i\gamma\,(\gamma + m_i)}{m_i} \int_0^\infty e^{-\gamma a}\left(1 - e^{-m_i a}\right)\big(1 - \alpha_i\tilde{n}_{j|0}(a)\big)da - (m_i + \gamma) > 0.$$

We then substitute the equilibrium density of species $j$ when it is on its own, i.e. $\tilde{n}_{j|0}(a) = \frac{r_j\tilde{S}_{j|0}}{m_j}\,(1 - e^{-m_j a})$.

$$\frac{1}{N_i}\frac{dN_i}{dt}\Big|_{\epsilon_i(a),\tilde{n}_{j|0}(a)} \approx \frac{r_i\gamma\,(\gamma + m_i)}{m_i} \int_0^\infty e^{-\gamma a}\left(1 - e^{-m_i a}\right)\Big(1 - \alpha_i\frac{r_j\tilde{S}_j}{m_j}\left(1 - e^{-m_j a}\right)\Big)da - (m_i + \gamma) > 0$$

Simplifying we have:

$$\frac{1}{N_i}\frac{dN_i}{dt}\Big|_{\epsilon_i(a),\tilde{n}_{j|0}(a)} \approx r_i - \frac{\alpha_i r_i r_j \tilde{S}_j\,(2\gamma + m_i + m_j)}{(\gamma + m_j)\,(\gamma + m_i + m_j)} - (m_i + \gamma) > 0.$$

$$\frac{1}{N_i}\frac{dN_i}{dt}\Big|_{\epsilon_i(a),\tilde{n}_{j|0}(a)} \approx r_i - \frac{\alpha_i r_i\,(2\gamma + m_i + m_j)\,(\gamma + 2m_j)\,(r_j - (\gamma + m_j))}{2\alpha_j r_j\,(\gamma + m_j)\,(\gamma + m_i + m_j)} - (m_i + \gamma) > 0.$$

Similarly, we have the condition for species $j$ to invade species $i$:

$$\frac{1}{N_j}\frac{dN_j}{dt}\Big|_{\tilde{n}_{i|0}(a),\epsilon_j(a)} \approx r_j - \frac{\alpha_j r_j\,(2\gamma + m_i + m_j)\,(\gamma + 2m_i)\,(r_i - (\gamma + m_i))}{2\alpha_i r_i\,(\gamma + m_i)\,(\gamma + m_i + m_j)} - (m_j + \gamma) > 0.$$

One can see these conditions are the same as the feasibility conditions (S4) and (S5).

#### 3.5 Local Stability

In the next two subsections we study the local stability of the equilibrium configurations for the single species and coexistence equilibrium. Assessing the linear equilibrium solution to an evolutionary PDE is akin to the stability analysis of an equilibrium in an ODE system. These methods in PDE are typical of the calculus of variations, see the classical [34, Chapter 8] or [33, Chapter 3] for a more gentle introduction.
Given $n^*$ an equilibrium for the dynamics, we will consider the linearized dynamics of a perturbation of the type

$$n = n^* + \epsilon\phi$$

where $\epsilon > 0$ is tiny and $\phi$ is a given perturbation that is square integrable with respect to the density of patches

$$\int_0^\infty \phi^2\rho\ da < \infty.$$

The patch distribution is the natural background measure to quantify the size of the perturbation. The choice of the $L^2$ avoids perturbations that are fully localized in a specific patch age, which could be argued are not ecologically relevant as the populations are assumed to disperse globally across patches and any perturbation affecting a given patch age would also affect the population across patches of different ages. Moreover, the choice of the weighed $L^2$ norm is extremely natural for the computations below.

##### 3.5.1 Single Species Equilibrium

Consider the PDE for one species at equilibrium on its own.

$$\frac{\partial n}{\partial t} = b\int_0^\infty \rho\ n(1-n)\ da - \mu n - \frac{\partial n}{\partial a}$$

We consider the dynamics of the following perturbation around the equilibrium $n^*$

$$n = n^* + \epsilon\phi,$$

which is given by

$$\begin{aligned}\frac{\partial n^*}{\partial t} + \epsilon\frac{\partial \phi}{\partial t} &= b\int_0^\infty \rho\ (n^*+\epsilon\phi)(1-(n^*+\epsilon\phi))\ da - \mu(n^*+\epsilon\phi) - \frac{\partial n^*}{\partial a} - \epsilon\frac{\partial\phi}{\partial a}\\ \epsilon\frac{\partial\phi}{\partial t} &= b\int_0^\infty \left(\epsilon\rho\phi - 2n^*\epsilon\rho\phi - \epsilon^2\rho\phi^2\right)\ da - \mu\epsilon\phi - \epsilon\frac{\partial\phi}{\partial a}.\end{aligned}$$

We divide both sides by $\epsilon$ and multiply both sides by $\phi$, and assume the $\epsilon^2$ terms are negligible.

$$\frac{\partial\phi}{\partial t}\phi = b\phi\int_0^\infty \left(\rho\phi(1-2n^*)\right)\ da - \mu\phi^2 - \frac{\partial\phi}{\partial a}\phi$$

We multiply both sides by $\rho$ and then integrate over patch-age $a$.

$$\frac{\partial}{\partial t}\int\phi^2\rho\ da = b\left(\int\phi\rho\ da\right)\left(\int_0^\infty \left(\rho\phi(1-2n^*)\right)\ da\right) - \mu\int\phi^2\rho\ da - \int\frac{\partial\phi}{\partial a}\phi\rho\ da$$

$$\frac{\partial}{\partial t}\int\ \phi^2\rho\ da = b\left(\int\ \phi\rho\ da\right)\left(\int_0^\infty \left(\rho\phi(1-2n^*)\right)\ da\right) - \left(\mu+\frac{\gamma}{2}\right)\int\ \phi^2\rho\ da.$$

We next apply the Cauchy-Schwarz inequality, (S9)

$$|\langle\mathbf{u},\mathbf{v}\rangle| \leq ||\mathbf{u}||\ ||\mathbf{v}||, \tag{S9}$$

multiple times to put a useful bound on the RHS of this dynamical equation. First, we use

$$|\langle\sqrt{\rho},\sqrt{\rho}\phi\rangle|_{L^2} \leq ||\sqrt{\rho}||_{L^2}||\sqrt{\rho}\phi||_{L^2}.$$

Resulting in

$$\int\rho\phi \leq \left(\int\rho\right)^{\frac{1}{2}}\left(\int\rho\phi^2\right)^{\frac{1}{2}}.$$

Since we know the integral of $\rho$ is 1 we have

$$\boxed{\int\rho\phi \leq \left(\int\rho\phi^2\right)^{\frac{1}{2}}.}$$

Thus we know

$$|\langle\sqrt{\rho}\phi,\sqrt{\rho}(1-2n^*)\rangle|_{L^2} \leq ||\sqrt{\rho}\phi||_{L^2}||\sqrt{\rho}(1-2n^*)||_{L^2}.$$

Resulting in

$$\boxed{\int\rho\phi(1-2n^*) \leq \left(\int\rho\phi^2\right)^{\frac{1}{2}}\left(\int\rho(1-2n^*)^2\right)^{\frac{1}{2}}.}$$

Together this results in

$$b\left(\int\ \phi\rho\ da\right)\left(\int_0^\infty \rho\phi(1-2n^*)\ da\right) \leq b\left(\int\rho\phi^2\ da\right)\left(\int_0^\infty \rho(1-2n^*)^2\ da\right)^{\frac{1}{2}}.$$

This results in

$$\begin{aligned}\frac{\partial}{\partial t}\int\ \phi^2\rho\ da &= \underbrace{b\left(\int\ \phi\rho\ da\right)\left(\int_0^\infty \left(\rho\phi(1-2n^*)^2\right)\ da\right)}_{\text{gain}} - \underbrace{\left(\mu+\frac{\gamma}{2}\right)\int\ \phi^2\rho\ da}_{\text{loss}}\\ \frac{\partial}{\partial t}\int\ \phi^2\rho\ da &\leq b\left(\int\rho\phi^2\ da\right)\left(\int_0^\infty \rho(1-2n^*)^2\ da\right)^{\frac{1}{2}} - \left(\mu+\frac{\gamma}{2}\right)\int\ \phi^2\rho\ da\\ &\leq \left(\int\rho\phi^2\ da\right)\left(b\left(\int_0^\infty \rho(1-2n^*)^2\ da\right)^{\frac{1}{2}} - \left(\mu+\frac{\gamma}{2}\right)\right).\end{aligned}$$

This results in in order for the equilibrium to be stable we need

$$\left(\mu + \frac{\gamma}{2}\right) \geq b \left(\int_0^\infty \rho \left(1 - 2n^*\right)^2 \, da\right)^{\frac{1}{2}}.$$

Inputting the density for one species on its own at equilibrium results in

$$\left(\mu + \frac{\gamma}{2}\right) \geq b \left(\int_0^\infty \rho \left(1 - 2\frac{(\gamma + 2\mu)\,(b - \gamma - \mu)}{2b\mu}\left(1 - e^{-\mu a}\right)\right)^2 da\right)^{\frac{1}{2}}$$

$$\left(\mu + \frac{\gamma}{2}\right) \geq \left(b^2 - 2b(\gamma + 2\mu) + 2(\gamma + \mu)(\gamma + 2\mu)\right)^{\frac{1}{2}}.$$

This is a sufficient but not necessary condition for local stability of the one species equilibrium. Numerical exploration of the condition indicates it is more stringent than the condition for feasibility of one species.

##### 3.5.2 Local Stability of Coexistence Equilibrium

Consider the PDE when both species are present at equilibrium.

$$\frac{\partial n_i}{\partial t} = r_i \int_0^\infty \rho \, n_i(1 - n_i - n_j) \, da - m_i n_i - \frac{\partial n_i}{\partial a}$$

We consider the perturbation (and the equivalent one for species $j$)

$$n_i = n_i^* + \epsilon\phi_i$$

resulting in

$$\frac{\partial n_i^*}{\partial t} + \epsilon\frac{\partial \phi_i}{\partial t} = r_i \int_0^\infty \rho \, \left(n_i^* + \epsilon\phi_i\right)\left(1 - \left(n_i^* + \epsilon\phi_i\right) - \left(n_j^* + \epsilon\phi_j\right)\right) \, da - m_i n_i^* - m_i\epsilon\phi_i - \frac{\partial n_i^*}{\partial a} - \epsilon\frac{\partial \phi_i}{\partial a}.$$

Expanding the terms in the integral and rearranging we have

$$\begin{aligned}\frac{\partial n_i^*}{\partial t} + \epsilon\frac{\partial \phi_i}{\partial t} =& - m_i n_i^* - m_i\epsilon\phi_i - \frac{\partial n_i^*}{\partial a} - \epsilon\frac{\partial \phi_i}{\partial a} + r_i \int_0^\infty \left(\rho n_i^* - \rho {n_i^*}^2 - \rho n_i^* n_j^*\right) \, da \\ &+ r_i \int_0^\infty \left(-\rho\epsilon\phi_i n_j^* - \rho\epsilon n_i^*\phi_j - \rho\epsilon^2\phi_i\phi_j - 2\rho\epsilon n_i^*\phi_i - \rho\epsilon^2\phi_i^2 + \rho\epsilon\phi_i\right) \, da.\end{aligned}$$

Again, similar to the one-species case we take any terms with $\epsilon^2$ to be negligible. This leaves us with

$$\epsilon\frac{\partial \phi_i}{\partial t} = \epsilon r_i \int_0^\infty \rho\left(-\phi_i n_j^* - n_i^*\phi_j - 2n_i^*\phi_i + \phi_i\right) \, da - m_i\epsilon\phi_i - \epsilon\frac{\partial \phi_i}{\partial a}.$$

We cancel an epsilon on both sides and multiply by either $\phi_i$ (or $\phi_j$ for species $j$.)

$$\frac{\partial \phi_i}{\partial t}\phi_i = r_i\phi_i \int_0^\infty \rho\left(-\phi_i n_j^* - n_i^*\phi_j - 2n_i^*\phi_i + \phi_i\right) \, da - m_i\phi_i^2 - \frac{\partial \phi_i}{\partial a}\phi_i$$

We multiply both sides by $\rho$ and then integrate both sides with respect to patch-age $a$.

$$\frac{\partial}{\partial t}\int \phi_i^2\rho \, da = r_i\left(\int \phi_i\rho \, da\right)\int_0^\infty \rho\left(\phi_i\left(1 - 2n_i^* - n_j^*\right) - \phi_j n_i^*\right) \, da - m_i\int \rho\phi_i^2 \, da - \int \frac{\partial \phi_i}{\partial a}\phi_i\rho \, da$$

We integrate the last term by parts in the same manner as the one species case, once again assuming $\phi_i(0)\rho(0) = \lim_{a\to\infty}\phi_i(a)\rho(a) = 0$ (similarly for species $j$) to get:

$$\frac{\partial}{\partial t}\int \phi_i^2\rho \, da = r_i\left(\int \phi_i\rho \, da\right)\int_0^\infty \rho\left(\phi_i\left(1 - 2n_i^* - n_j^*\right) - \phi_j n_i^*\right) \, da - \left(m_i + \frac{\gamma}{2}\right)\int \rho\phi_i^2 \, da.$$

We separate the parts of these equations which still depend on the $\phi_i$ and $\phi_j$ functions.

$$\begin{aligned}\frac{\partial}{\partial t}\int \phi_i^2\rho \, da =& r_i\left(\int \phi_i\rho \, da\right)\int_0^\infty \rho\phi_i\left(1 - 2n_i^* - n_j^*\right) \, da - r_i\left(\int \phi_i\rho \, da\right)\int_0^\infty \rho\phi_j n_i^* \, da \\ &- \left(m_i + \frac{\gamma}{2}\right)\int \rho\phi_i^2 \, da.\end{aligned} \tag{S10}$$

We next apply the Cauchy-Schwarz ((S9)) inequality multiple times to put useful bounds on the RHS of this dynamic equation. First we use

$$|\langle\sqrt{\rho}, \sqrt{\rho}\phi_i\rangle|_{L^2} \leq ||\sqrt{\rho}||_{L^2}||\sqrt{\rho}\phi_i||_{L^2}.$$

resulting in

$$\int \rho\phi_i \leq \left(\int \rho\right)^{\frac{1}{2}}\left(\int \rho\phi_i^2\right)^{\frac{1}{2}}.$$

Since we know the integral of $\rho$ over all patch ages is 1 we have

$$\int \rho\phi_i \leq \left(\int \rho\phi_i^2\right)^{\frac{1}{2}} \tag{S11}$$

We have a similar inequality for $\phi_j$ and $\rho$.Thus we know, again using Cauchy Schwarz, that

$$|\langle\sqrt{\rho}\phi_i, \sqrt{\rho}\left(1-2n_i^*-n_j^*\right)\rangle \leq ||\sqrt{\rho}\phi_i||_{L^2}||\sqrt{\rho}\left(1-2n_i^*-n_j^*\right)||_{L^2}.$$

Which results in

$$\int \rho\phi_i(1-2n_i^*-n_j^*) \leq \left(\int \rho\phi_i^2\right)^{\frac{1}{2}}\left(\int_0^\infty \rho\left(1-2n_i^*-n_j^*\right)^2\right)^{\frac{1}{2}}. \tag{S12}$$

Again, using Cauchy Schwarz, we have

$$|\langle\sqrt{\rho}\phi_j, \sqrt{\rho}n_i^*\rangle \leq ||\sqrt{\rho}\phi_j||_{L^2}||\sqrt{\rho}n_i^*||_{L^2}$$

$$\int \rho\phi_j n_i^* \leq \left(\int \rho\phi_j^2\right)^{\frac{1}{2}}\left(\int_0^\infty \rho n_i^{*2}\right)^{\frac{1}{2}}.$$

Together (S11) and (S12) tell us

$$r_i\left(\int \phi_i\rho\right)\int_0^\infty \rho\phi_i\left(1-2n_i^*-n_j^*\right) \leq r_i\left(\int \rho\phi_i^2\right)\left(\int_0^\infty \rho\left(1-2n_i^*-n_j^*\right)^2\right)^{\frac{1}{2}}. \tag{S13}$$

Using (S13) with (S10) which results in

$$\frac{\partial}{\partial t}\int \phi_i^2\rho \leq \left(\int_0^\infty \rho\phi_i^2\right)\left[r_i\left(\int_0^\infty \rho\left(1-2n_i^*-n_j^*\right)^2\right)^{\frac{1}{2}} - \left(m_i+\frac{\gamma}{2}\right)\right] - r_i\int \phi_i\rho\left(\int_0^\infty \rho\phi_j n_i^*\right). \tag{S14}$$

Again by Cauchy Schwarz we know

$$|\langle\sqrt{\rho}\phi_j, \sqrt{\rho}n_i^*\rangle| \leq ||\sqrt{\rho}\phi_j||_{L^2}||\sqrt{\rho}n_i^*||_{L^2}.$$

This means we know

$$\int \phi_j\rho n_i^* \leq \left(\int \rho\phi_j^2\right)^{\frac{1}{2}}\left(\int \rho n_i^{*2}\right)^{\frac{1}{2}}. \tag{S15}$$

Using (S14) with (S15) which results in

$$\frac{\partial}{\partial t}\int \phi_i^2\rho \leq \left(\int_0^\infty \rho\phi_i^2\right)\left[r_i\left(\int_0^\infty \rho\left(1-2n_i^*-n_j^*\right)^2\right)^{\frac{1}{2}} - \left(m_i+\frac{\gamma}{2}\right)\right] - r_i\left(\int \rho\phi_i^2\right)^{\frac{1}{2}}\left(\int \rho\phi_j^2\right)^{\frac{1}{2}}\left(\int \rho n_i^{*2}\right)^{\frac{1}{2}}.$$

We can rewrite this system as

$$\frac{\partial}{\partial t}X \leq X\left[r_i\left(\int_0^\infty \rho\left(1-2n_i^*-n_j^*\right)^2\right)^{\frac{1}{2}} - \left(m_i+\frac{\gamma}{2}\right)\right] - r_i\left(X\right)^{\frac{1}{2}}\left(Y\right)^{\frac{1}{2}}\left(\int \rho n_i^{*2}\right)^{\frac{1}{2}}.$$

Where

$$X = \int_0^\infty \rho\phi_i^2\, da$$
$$Y = \int_0^\infty \rho\phi_j^2\, da.$$

This time we use the arithmetic-geometric mean inequality to simplify, resulting in :

$$\frac{\partial}{\partial t}X \leq X\left[r_i\left(\int_0^\infty \rho\left(1-2n_i^*-n_j^*\right)^2\right)^{\frac{1}{2}} - \left(m_i+\frac{\gamma}{2}\right)\right] - \frac{r_i}{2}\left[X+Y\right]\left(\int \rho n_i^{*2}\right)^{\frac{1}{2}}.$$

We further simplify to get

$$\frac{\partial}{\partial t}X \leq X\left[r_i\left(\int_0^\infty \rho\left(1-2n_i^*-n_j^*\right)^2\right)^{\frac{1}{2}} - \left(m_i+\frac{\gamma}{2}\right) - \frac{r_i}{2}\left(\int \rho n_i^{*2}\right)^{\frac{1}{2}}\right] - \frac{r_i}{2}\left(\int \rho n_i^{*2}\right)^{\frac{1}{2}}Y.$$

The work for species $j$ is done in a similar fashion.

This results in the system

$$\frac{\partial}{\partial t}\begin{bmatrix} X \\ Y \end{bmatrix} \leq \mathbf{A}\begin{bmatrix} X \\ Y \end{bmatrix}$$

where

$$\mathbf{A} = \begin{bmatrix} r_i\left(\int_0^\infty \rho\left(1-2n_i^*-n_j^*\right)^2\, da\right)^{\frac{1}{2}} - (m_i+\frac{\gamma}{2}) - \frac{r_i}{2}\left(\int \rho n_i^{*2}\, da\right)^{\frac{1}{2}} - \frac{r_i}{2}\left(\int \rho n_i^{*2}\, da\right)^{\frac{1}{2}} & r_j\left(\int_0^\infty \rho\left(1-n_i^*-2n_j^*\right)^2\, da\right)^{\frac{1}{2}} - (m_j+\frac{\gamma}{2}) - \frac{r_j}{2}\left(\int \rho n_j^{*2}\, da\right)^{\frac{1}{2}} - \frac{r_j}{2}\left(\int \rho n_j^{*2}\, da\right)^{\frac{1}{2}} \end{bmatrix}$$

Hence sufficient but not necessary conditions for local stability is that the dominant eigenvalue of the matrix $\mathbf{A}$ must have a negative real part. The matrix and its eigenvalues can be calculated in terms of the model parameters but the resulting expressions are quite complicated. Numerical exploration of the condition that the dominant eigenvalue has a negative real part indicates that this condition holds whenever the two species equilibrium is feasible.

#### 3.6 Coexistence or Not from Species Variation on Trade-Offs

Here we will show that our coexistence conditions for this model case can be satisfied if species $i$ and $j$ vary appropriately along demographic trade-offs. Our conditions for feasibility of coexistence and mutual invasion can be summarized as

$$\frac{(\gamma+2m_i)(2\gamma+m_i+m_j)}{2(\gamma+m_i)(\gamma+m_i+m_j)} < \frac{\frac{r_i\alpha_i}{(r_i-\gamma-m_i)}}{\frac{r_j\alpha_j}{(r_j-\gamma-m_j)}} < \frac{2(\gamma+m_j)(\gamma+m_i+m_j)}{(\gamma+2m_j)(2\gamma+m_i+m_j)} \tag{S16}$$

with the right inequality being the condition for feasibility of species $i$ (or for it to invade species $j$) and the left inequality being the condition for feasibility of species $j$ (or for it to invade species $i$). See Fig. 1 for a comparison between analytical ((S16)) and numerical coexistence regions, Fig. 2 for a look at how the coexistence region varies when disturbance changes.

We consider a variety of trade-offs throughout the different model cases and have listed them all, and the results we obtained for each, in Table 1.

##### 3.6.1 Coexistence from Reproduction/Adult-Survival Trade-Off

We first consider whether variation between species on a reproduction/adult-survival trade-off can enable this series of inequalities to hold. In other words, we will consider whether if $m_i > m_j$ (species $j$ has better survival of its adults), and species are equal in their other parameters ($\alpha_i = \alpha_j$), does there always exist an $r_i > r_j$ (making species $i$ the better reproducer) that will enable the series of inequalities in (S16) to hold?

It turns out one can show (we leave this as an exercise to the reader) that if one takes $m_i = m_j + \Delta_m$ and $r_i = r_j + \Delta_r$ that (S16) implies (so long as species $i$ and $j$ are able to persist on their own and $r_j$ is not *too* large) that

$$\frac{r_j\Delta_m\left(4m_j^2+7m_j\gamma-r_j\gamma+3\gamma^2+2\left(m_j+\gamma\right)\Delta_m\right)}{\left(2m_j+\gamma\right)^2\left(2m_j+\gamma\right)+\left(2m_j^2+3m_j\gamma+\gamma\left(r_j+\gamma\right)\right)\Delta_m} < \Delta_r < \frac{r_j\Delta_m\left(4m_j^2+7m_j\gamma-r_j\gamma+3\gamma^2+2\left(m_j+\gamma\right)\Delta_m\right)}{2\left(m_j+\gamma\right)^2\left(2m_j+\gamma\right)+\left(6m_j^2+10m_j\gamma+\gamma\left(r_j+4\gamma\right)\right)\Delta_m+2\left(m_j+\gamma\right)\Delta_m^2}. \tag{S17}$$

The specific limitation on $r_j$ for this series of inequalities to apply is

$$r_j < \frac{1}{\gamma}\left(4m_j^2+7m_j\gamma+3\gamma^2+2\left(m_j+\gamma\right)\Delta_m\right).$$

Our analytical calculation of (S16) is unlikely to apply outside of this regime anyway, since higher values of $r_j$ will make species $j$'s density higher and more likely to make the linear density-dependent term go to zero. Note that in (S17), the left inequality is now the one having to do with the ability of species $i$ to invade species $j$, and hence it is telling us what is the minimum advantage $\Delta_r$ species $i$ must have in reproduction to overcome its disadvantage in survival ($\Delta_m$) to coexist. The right inequality is telling us the conditions for species $j$ to be able to invade species $i$ and, as expected, is setting an upper limit on the advantage species $i$ can have in reproduction while still allowing species $j$ to invade it. Also note that the competition sensitivities $\alpha_i = \alpha_j = \alpha$ simply divide out of the mutual invasion inequality, since they only appear in the middle term as a ratio. So the overall magnitude of the competition sensitivity has no influence here.

In the regime of $r_j$ in which the inequality holds, we show that the RHS of (S17) is larger than the LHS for all $\Delta_m$, so long as species $j$ can persist on its own (requiring $r_j > m_j + \gamma$). We also note the right and left most sides of (S17) are both positive, indicating that a positive $\Delta_r$ is needed (as expected), and also that $\lim_{\Delta_m \to 0} \Delta_r = 0$.

Hence we have shown that for any $m_i > m_j$ (any $\Delta_m > 0$) there exists a range of $r_i > r_j$ (of $\Delta_r > 0$) such that species $i$ and $j$ can have feasible coexistence with mutual invasion, so variation between species $i$ and $j$ on a reproduction/adult-survival trade-off will enable them to coexist.

Furthermore, for an arbitrarily small $\Delta_m$ this will still be true, so there will be no inherent limit to the similarity of the two species that are coexisting, so long as both $r_i$ and $m_i$ are free to vary independently, then $\Delta_r$ can satisfy the inequality (S17).

##### 3.6.2 Coexistence from Adult-Survival-Robustness to Competition Trade-Off

We next consider whether variation between species on an adult-survival versus robustness to competition trade-off can enable this series of inequalities to hold. Here the kind of "robustness to competition" we look at is the robustness of reproduction to competition, since that is the component of demography upon which competition is acting on in this model case.

Specifically, we will consider whether if $m_i < m_j$ (making species $i$ the better survivor), when species are equal in their other parameters ($r_i = r_j = r$), does there always exists an $\alpha_i > \alpha_j$ (making species $j$ have greater "robustness to competition", i.e. lower sensitivity $\alpha$) that will enable this series of inequalities in (S16) to hold (presuming other parameters are equal across the species)?

It turns out one can show (the algebra is messy so we will not bore the reader with the details) that if one takes $m_i = m_j - \Delta_m$ and $\alpha_i = \alpha_j + \Delta_\alpha$ (S16) implies (so long as species $i$ and $j$ can persist on their own and $r_i = r_j = r$ is not *too* large) that

$$\frac{\Delta_m \alpha_j \left(3\gamma^2 - 5\gamma\Delta_m + 7\gamma m_j - 6\Delta_m m_j + 4m_j^2 - \gamma r + 2\Delta_m^2\right)}{2\left(\gamma + m_j - \Delta_m\right)\left(\gamma + 2m_j - \Delta_m\right)\left(-\gamma - m_j + r\right)} < \Delta_\alpha < \frac{\Delta_m \alpha_j \left(3\gamma^2 - 2\Delta_m\left(\gamma + m_j\right) + 7\gamma m_j + 4m_j^2 - \gamma r\right)}{\left(\gamma + 2m_j\right)\left(2\gamma + 2m_j - \Delta_m\right)\left(-\gamma - m_j + r\right)}.$$

Again, our analytical calculation of (S16) is unlikely to hold outside of this regime. As in Sec. 3.6.1 the left inequality represents the ability of species $i$ to invade species $j$. The right inequality represents the ability of species $j$ to invade species $i$.

Once more we find both the LHS and the RHS of our inequalities hold positive and thus a positive $\Delta_\alpha$ is needed (as expected). We are able to show that the RHS of the inequality is greater than the LHS of the inequality as long as the species are able to persist on their own. See Fig. 3 for a comparison between analytical and numerical coexistence regions and Fig. 4 for a look at how the coexistence region changes as we vary disturbance.

Hence, we have shown that for any $m_i < m_j$ (any $\Delta_m > 0$) there exists a range of competition sensitivities of species $i$ ($\alpha_i > \alpha_j$ or $\Delta_\alpha$) such that the two species can have feasible coexistence with mutual invasion. This trade-off between adult-survival and robustness of reproduction to competition allows for the two species to coexist, and there is no particular limit to the similarity of the potentially coexisting species in this 2-species case.

##### 3.6.3 *No* Coexistence from Reproduction-Robustness of Reproduction to Competition Trade-Off

Finally, it is straighforward to see that the series of inequalities in (S16) *cannot* be satisfied by variation between species in reproduction and robustness of reproduction to competition alone, i.e. by $r_i \neq r_j$ and $\alpha_i \neq \alpha_j$. This is because both of these parameters influence *only the middle term* of (S16), and when $m_i = m_j$ the most RHS of the series of inequalities is equal to the most LHS. So there is no room for the middle term to fit.

#### 3.7 Which of the Coexisting Species Would be the Better Competitor for a Single Patch when Disturbance is removed

We constructed our model such that without patch dynamics it would not yield competitive coexistence (see Section 2). Here we ask, if two species coexist in this model case through a reproduction/adult-survival trade-off, which one, the higher reproducing species, or the better survivor, would win in competition in a single isolated patch with disturbance and patch dynamics, and in this sense be the "better within-patch competitor"? This question is relevant not only as a curiosity, but to help compare this coexistence-enabling trade-off with the well-known "competition-colonization" trade-off, in which a better colonizer of new patches (presumably the better reproducer, as it is assumed to have a faster rate of colonization of vacant patches by propagules coming from patches it occupies) can coexist with a species that is the better competitor for patches (it can take over patches from the better colonizer).

Let's assume species $i$ has $r_i > r_j$ and $m_i > m_j$, i.e. it is the higher reproducer while species $j$ is the better survivor. For species $i$ to be the *better competitor*, i.e. the winner without patch dynamics and disturbance in this model case (i.e. for it to invade species $j$ but not vice versa since it would require the opposite condition), from Section 2. Outlining the dynamics our model would have *without disturbance* we see we need:

$$\frac{r_i\alpha_i}{(r_i - m_i)} < \frac{r_j\alpha_j}{(r_j - m_j)}.$$

One can show by plugging in $m_i = m_j + \Delta_m$ and $r_i = r_j + \Delta_r$ that this implies

$$\frac{\Delta_r}{r_j} > \frac{\Delta_m}{m_j}.$$

We can check whether this holds when there is coexistence of the two species under disturbance and patch dynamics. First, let us consider the LHS of the inequality (S17) which is the criterion for species $i$ to invade species $j$:

$$\frac{\Delta_m\left(3\gamma^2 + 2\Delta_m\left(\gamma + m_j\right) + 7\gamma m_j + 4m_j^2 - \gamma r_j\right)}{2\left(\gamma + 2m_j\right)\left(\gamma + m_j\right)^2 + \Delta_m\left(3\gamma m_j + 2m_j^2 + \gamma\left(\gamma + r_j\right)\right)} < \frac{\Delta_r}{r_j}.$$

So we want to know if, given this holds, would it imply that species $i$ is the better competitor? This would require

$$\frac{\Delta_m}{m_j} < \frac{\Delta_m\left(3\gamma^2 + 2\Delta_m\left(\gamma + m_j\right) + 7\gamma m_j + 4m_j^2 - \gamma r_j\right)}{2\left(\gamma + 2m_j\right)\left(\gamma + m_j\right)^2 + \Delta_m\left(3\gamma m_j + 2m_j^2 + \gamma\left(\gamma + r_j\right)\right)}.$$

We can show this can hold sometimes, though in a somewhat limited region of parameter space from numerical exploration.

However, for mutual invasion to occur, we also need species $j$ to be able to invade species $i$. This is determined by the other half of (S17), that provides an upper constraint on $\Delta_r$. Dividing both sides of that part of the inequality by $r_j$ we see that what we need:

$$\frac{\Delta_m}{m_j} < \frac{\Delta_r}{r_j} < \frac{\Delta_m\left((\gamma+\Delta_m)(3\gamma+2\Delta_m)+(7\gamma+6\Delta_m)m_j+4m_j^2-\gamma r_j\right)}{2\Delta_m^2\left(\gamma+m_j\right)+2\left(\gamma+m_j\right)^2\left(\gamma+2m_j\right)+\Delta_m\left(10\gamma m_j+6m_j^2+\gamma\left(4\gamma+r_j\right)\right)}.$$

We check whether this is satisfied, i.e. whether

$$\frac{\Delta_m}{m_j} < \frac{\Delta_m\left((\gamma+\Delta_m)(3\gamma+2\Delta_m)+(7\gamma+6\Delta_m)m_j+4m_j^2-\gamma r_j\right)}{2\Delta_m^2\left(\gamma+m_j\right)+2\left(\gamma+m_j\right)^2\left(\gamma+2m_j\right)+\Delta_m\left(10\gamma m_j+6m_j^2+\gamma\left(4\gamma+r_j\right)\right)}$$

is true, and we find it is not.

Hence when species $i$ and $j$ coexist through a reproduction/adult-survival trade-off under disturbance (with species $i$ being the better reproducer) it is *never* the better competitor in the absence of disturbance. In this sense the trade-off has some similarity with the competition-colonization trade-off, in that species $i$, being the better reproducer, could be viewed as the better "colonizer" and species $j$, the one with higher adult-survival, as the better competitor. However, the differences are indeed just in basic demographic rate rather than sensitivity to competition, and in this sense the trade-off is a bit different. Also, our reproduction-survival trade-off does not involve exclusion of species $i$ from patches as they age when species $j$ is also there. Instead, in our model, both species remain present at old patch-ages, because propagules produced in other patches keep species $i$ present in older patches.

#### 3.8 Non-Dimensionalizing

We non-dimensionalize the system and in the process reduce the number of model parameters.

First we must assume we are in the regime of a non-existent critical patch-age cut-off, i.e. that there are no non-negative and finite values of $a$ which satisfy

$$0 = 1-\alpha_i n_i(a) \qquad \text{or} \qquad 0 = 1-\alpha_i n_i(a)-\alpha_i n_j(a)$$

or the corresponding species $j$ equations.

Then we consider the original equations for the densities of both species:

$$\frac{\partial n_i}{\partial t} = r_i\int_0^\infty \rho(a',t)n_i(a',t)\big(1-\alpha_i n_i(a',t)-\alpha_i n_j(a',t)\big)da' - m_i n_i(a,t) - \frac{\partial n_i(a,t)}{\partial a}$$
$$\frac{\partial n_j}{\partial t} = r_j\int_0^\infty \rho(a',t)n_j(a',t)\big(1-\alpha_j n_i(a',t)-\alpha_j n_j(a',t)\big)da' - m_j n_j(a,t) - \frac{\partial n_j(a,t)}{\partial a}$$

$$\frac{\rho(a,t)}{\partial t} = -\frac{\partial\rho(a,t)}{\partial a} - \gamma\rho(a,t)$$
$$\rho(0,t) = \int_0^\infty \gamma\rho(a,t)\;da$$
$$n_i(0) = 0.$$

We have the following **parameters:** $\gamma, \alpha_i, \alpha_j r_i, r_j, m_i, m_j$.
We have the following **variables:** $n_i, n_j, \rho, a, t$.
The *goal* is the convert $n_i, n_j, \rho, a, t$ to the dimensionless $\hat{n}_i, \hat{n}_j, \hat{\rho}, \hat{a}, \hat{t}$ in terms of $\gamma, \alpha_i, \alpha_j, r_i, r_j, m_i, m_j$.
The *technique*: we set

$$\hat{n}_i = \alpha_i\frac{n_i-n_{ir}}{n_{i0}}, \quad \hat{n}_j = \alpha_j\frac{n_j-n_{jr}}{n_{j0}}, \quad \hat{\rho} = \frac{\rho-\rho_r}{\rho_0}, \quad \hat{a} = \frac{a-a_r}{a_0}, \quad \hat{t} = \frac{t-t_r}{t_0}.$$

Letting $n_{ir}, n_{jr}, \rho_r, a_r, t_r$ be the reference values, and $n_{i0}, n_{j0}, \rho_0, a_0, t_0$ be the scaling factors.

We assume $n_i, n_j, \rho, a, t$ begin at $n_i(0)=0, n_j(0)=0, \rho(0)=0, a(0)=0, t(0)=0$.
This results in $n_{ir}=n_{jr}=\rho_r=a_r=t_r=0$.

This simplifies the nondimensional variables and time to

$$\hat{n}_i = \alpha_i\frac{n_i}{n_{i0}}, \quad \hat{n}_j = \alpha_j\frac{n_j}{n_{j0}}, \quad \hat{\rho} = \frac{\rho}{\rho_0}, \quad \hat{a} = \frac{a}{a_0}, \quad \hat{t} = \frac{t}{t_0}.$$

We substitute $\frac{\hat{n}_i n_{i0}}{\alpha_i} = n_i$, $\frac{\hat{n}_j n_{j0}}{\alpha_j} = n_j$, $\hat{\rho}\rho_0 = \rho$, $\hat{a}a_0 = a$, $\hat{t}t_0 = t$

$$\frac{\partial\rho}{\partial t} = -\frac{\partial\rho}{\partial a} - \gamma\rho$$
$$\frac{\partial\hat{\rho}\cancel{\rho_0}}{\partial\hat{t}t_0} = -\frac{\partial\hat{\rho}\cancel{\rho_0}}{\partial\hat{a}a_0} - \gamma\hat{\rho}\cancel{\rho_0}$$
$$\frac{1}{t_0}\frac{\partial\hat{\rho}}{\partial\hat{t}} = -\frac{1}{a_0}\frac{\partial\hat{\rho}}{\partial\hat{a}} - \gamma\hat{\rho}.$$

We choose

$$t_0 = a_0 = \frac{1}{\gamma}\ \text{time}^{-1}.$$

Resulting in

$$\frac{\partial \hat{\rho}}{\partial \hat{t}} = -\frac{\partial \hat{\rho}}{\partial \hat{a}} - \hat{\rho}$$

$$\begin{aligned}
\rho(0,t) &= \int_0^\infty \gamma\rho\ da \\
\hat{\rho}\cancel{\rho_0}(0,t) &= \int_0^\infty \gamma\hat{\rho}\cancel{\rho_0}\ d\hat{a}a_0 \\
\hat{\rho}(0,t) &= \int_0^\infty \hat{\rho}\ d\hat{a}.
\end{aligned}$$

Together this results in

$$\hat{\rho}(a) = e^{-\hat{a}}.$$

The species equations (rewritten without substitution) are:

$$\begin{aligned}
\frac{\partial n_i}{\partial t} &= r_i \int_0^\infty \rho\ n_i\big(1 - \alpha_i n_i - \alpha_i n_j\big)da' - m_i n_i - \frac{\partial n_i}{\partial a} \\
\frac{\partial n_j}{\partial t} &= r_j \int_0^\infty \rho\ n_j\big(1 - \alpha_j n_i - \alpha_j n_j\big)da' - m_j n_j - \frac{\partial n_j}{\partial a}.
\end{aligned}$$

Applying the substitutions, simplifying and grouping the following

$$\frac{r_i}{\gamma} = \beta_i, \qquad \frac{r_j}{\gamma} = \beta_j, \qquad \frac{m_i}{\gamma} = \delta_i, \qquad \frac{m_j}{\gamma} = \delta_j, \qquad \frac{\alpha_i}{\alpha_j} = \kappa.$$

this results in

$$\begin{aligned}
\frac{\partial \hat{n}_i}{\partial \hat{t}} &= \beta_i \int_0^\infty \hat{\rho}\ \hat{n}_i\big(1 - \hat{n}_i - \kappa\hat{n}_j\big)d\hat{a}' - \delta_i\hat{n}_i - \frac{\partial \hat{n}_i}{\partial \hat{a}} \\
\frac{\partial \hat{n}_j}{\partial \hat{t}} &= \beta_j \int_0^\infty \hat{\rho}\ \hat{n}_j\big(1 - \frac{1}{\kappa}\hat{n}_i - \hat{n}_j\big)d\hat{a}' - \delta_j\hat{n}_j - \frac{\partial \hat{n}_j}{\partial \hat{a}}.
\end{aligned}$$

Finally we find:

$$\begin{aligned}
n_i(0) &= 0 \\
n_{i0}\hat{n}_i(0) &= 0 \\
\hat{n}_i(0) &= 0.
\end{aligned}$$

Resulting in

$$\begin{aligned}
\frac{\partial \hat{n}_i}{\partial \hat{t}} &= \beta_i \int_0^\infty \hat{\rho}\ \hat{n}_i\big(1 - \hat{n}_i - \kappa\hat{n}_j\big)d\hat{a}' - \delta_i\hat{n}_i - \frac{\partial \hat{n}_i}{\partial \hat{a}} \\
\frac{\partial \hat{n}_j}{\partial \hat{t}} &= \beta_j \int_0^\infty \hat{\rho}\ \hat{n}_j\big(1 - \frac{1}{\kappa}\hat{n}_i - \hat{n}_j\big)d\hat{a}' - \delta_j\hat{n}_j - \frac{\partial \hat{n}_j}{\partial \hat{a}} \\
\frac{\partial \hat{\rho}}{\partial \hat{t}} &= -\frac{\partial \hat{\rho}}{\partial \hat{a}} - \hat{\rho} \\
\hat{\rho} &= \int_0^\infty \hat{\rho}\ d\hat{a} \\
\hat{n}_i(0) &= 0.
\end{aligned}$$

Hence we end up with 4 new dimensionless variables, a dimensionless definition of time, and 5 parameters—2 fewer parameters than we started with. These parameters were absorbed either into our dimensionless variables, into our definitions of the dimensionless species densities or into our dimensionless time.

Note this tells us that all model behavior, including coexistence, is determined only by the values of dimensionless parameters $\frac{r_i}{\gamma}$, $\frac{r_j}{\gamma}$ and $\frac{m_i}{\gamma}$, $\frac{m_j}{\gamma}$, and the ratio $\frac{\alpha_i}{\alpha_j}$, and *not* the disturbance rate $\gamma$ in itself. This should also be true for our other model cases. However we chose not to plot results in this scaled parameter space because it clouds the dependence of the coexistence regime on the disturbance rate $\gamma$. The mapping between coexistence regions in scaled parameter space and real parameter space for different disturbance rates is complex. Not only will the scale of the axes in real parameter space vary with disturbance rate $\gamma$, but the scaled demographic strategy with which mutual invasion is being considered will also vary.

### 4 Competition Acting on Offspring-Survival

The second model case we consider has negative density-dependence on offspring survival and a density-independent reproduction/mortality. In the competition acting on offspring-survival case the chances of the offspring establishing themselves and growing to maturity depend on the adult density in the patch. The model equations for this case are

$$\frac{\partial n_i(a,t)}{\partial t} = \underbrace{r_i\Big(\int_0^\infty \rho(a',t)\; n_i(a',t)\Big)da' \max[0,(1-\alpha_i n_i(a,t)-\alpha_i n_j(a,t))]\; da'}_{\text{recruitment}} - \underbrace{m_i n_i(a,t)}_{\text{mortality}} - \underbrace{\frac{\partial n_i(a,t)}{\partial a}}_{\text{patch-aging}} \tag{S18}$$

$$\frac{\partial \rho(a,t)}{\partial t} = -\frac{\partial \rho(a,t)}{\partial a} - \gamma\rho(a,t)$$

$$\rho(0,t) = \int_0^\infty \gamma\rho(a,t)da$$

$$\text{at equilibrium in a patch of age } a=0, \quad n_i(0)=0.$$

#### 4.1 Obtaining Dynamical Equations for the Average Landscape Density

We were able to show (for cases where species' densities remain small enough that they do not drive the density-dependent factor in the recruitment term to 0) that the dynamical equations for the average landscape density of the two species in this model are equivalent to the Lotka-Volterra competition model. First we identify the average landscape density already present in the partial differential equation (PDE) for $n_i(a,t)$:

$$\frac{\partial n_i(a,t)}{\partial t} = r_i \underbrace{\Big(\int_0^\infty \rho(a',t)\; n_i(a',t)\Big)da'}_{N_i(t)} \max[0,(1-\alpha_i n_i(a,t)-\alpha_i n_j(a,t))] - m_i n_i(a,t) - \frac{\partial n_i(a,t)}{\partial a}.$$

Hence we have:

$$\frac{\partial n_i(a,t)}{\partial t} = r_i N_i(t)\max[0,(1-\alpha_i n_i(a,t-\alpha_i n_j(a,t)))] - m_i n_i(a,t) - \frac{\partial n_i(a,t)}{\partial a}.$$

We then define $\frac{\partial g_i(a,t)}{\partial t}$ (where $g_i(a,t) = n_i(a,t)\rho(a,t)$) and consider its time derivative, in order to obtain PDEs that can be integrated over patch-ages to get an ordinary differential equations (ODE) for the average landscape densities:

$$\frac{\partial g_i(a,t)}{\partial t} = \frac{\partial n_i(a,t)}{\partial t}\rho(a,t) + n_i(a,t)\frac{\partial \rho(a,t)}{\partial t}$$

$$\frac{\partial g_i(a,t)}{\partial t} = r_i N_i(t)\max[0,(1-\alpha_i n_i(a,t)-\alpha_i n_j(a,t))]\rho(a,t) - m_i \underbrace{n_i(a,t)\rho(a,t)}_{g_i(a,t)} - \frac{\partial n_i(a,t)}{\partial a}\rho(a,t) + n_i(a,t)\underbrace{\frac{\partial \rho(a,t)}{\partial t}}_{-\frac{\partial\rho(a,t)}{\partial a}-\gamma\rho(a,t)}$$

$$\begin{aligned}\frac{\partial g_i(a,t)}{\partial t} &= r_i N_i(t)\max[0,(1-\alpha_i n_i(a,t)-\alpha_i n_j(a,t))]\rho(a,t) - m_i g_i(a,t) - \frac{\partial n_i(a,t)}{\partial a}\rho(a,t) \\ &\quad + n_i(a,t)\left(-\frac{\partial \rho(a,t)}{\partial a} - \gamma\rho(a,t)\right) \\ \frac{\partial g_i(a,t)}{\partial t} &= r_i N_i(t)\max[0,(1-\alpha_i n_i(a,t)-\alpha_i n_j(a,t))]\rho(a,t) - m_i g_i(a,t) \\ &\quad - \underbrace{\left(\frac{\partial n_i(a,t)}{\partial a}\rho(a,t) + \frac{\partial \rho(a,t)}{\partial a}n_i(a,t)\right)}_{\frac{\partial g_i(a,t)}{\partial a}} - \gamma\underbrace{n_i(a,t)\rho(a,t)}_{g_i(a,t)}\end{aligned}$$

$$\frac{\partial g_i(a,t)}{\partial t} = r_i N_i(t)\max[0,(1-\alpha_i n_i(a,t)-\alpha_i n_j(a,t))]\rho(a,t) - (\gamma+m_i)\,g_i(a,t) - \frac{\partial g_i(a,t)}{\partial a}.$$

We next integrate both sides with respect to $a$.

$$\begin{aligned}\underbrace{\int_0^\infty da\frac{\partial g_i(a,t)}{\partial t}}_{\frac{dN_i}{dt}} &= \int_0^\infty da\; r_i N_i(t)\max[0,(1-\alpha_i n_i(a,t)-\alpha_i n_j(a,t))]\rho(a,t) - (\gamma+m_i)\underbrace{\int_0^\infty da\; g_i(a,t)}_{N_i} - \int_0^\infty da\frac{\partial g_i(a,t)}{\partial a} \\ \frac{dN_i}{dt} &= \int_0^\infty da\; r_i N_i(t)\max[0,(1-\alpha_i n_i(a,t)-\alpha_i n_j(a,t))]\rho(a,t) - (\gamma+m_i)\,N_i - \underbrace{g_i(\infty,t)-g_i(0,t)}_{=0} \\ \frac{dN_i}{dt} &= r_i N_i\int_0^\infty da\;\max[0,(1-\alpha_i n_i(a,t)-\alpha_i n_j(a,t))]\rho(a,t) - (\gamma+m_i)\,N_i.\end{aligned}$$

If instead the linear density-dependent term never crosses zero (because species densities simply don't get high enough in the patch), then

$$\begin{aligned}\frac{dN_i}{dt} &= r_i N_i(t)\int_0^\infty da\;(1-\alpha_i n_i(a,t)-\alpha_i n_j(a,t))\rho(a,t) - (\gamma+m_i)\,N_i(t) \\ \frac{dN_i}{dt} &= r_i N_i(t)\,(1-\alpha_i N_i(t)-\alpha_i N_j(a,t)) - (\gamma+m_i)\,N_i(t).\end{aligned}$$

This can be rearranged into the classical form of the Lotka-Voterra competition model, which for both species $i$ and $j$ is

$$\frac{dN_i}{dt} = \left(r_i - (\gamma + m_i)\right) N_i(t) \left(1 - \frac{r_i\alpha_i}{r_i - (\gamma + m_i)} N_i(t) - \frac{r_i\alpha_i}{r_i - (\gamma + m_i)} N_j(a,t)\right)$$
$$\frac{dN_j}{dt} = \left(r_j - (\gamma + m_j)\right) N_i(t) \left(1 - \frac{r_j\alpha_j}{r_j - (\gamma + m_j)} N_i(t) - \frac{r_j\alpha_j}{r_j - (\gamma + m_j)} N_j(a,t)\right).$$

Invasion analysis is straightforward in the Lotka-Volterra competition model. For species $i$ to invade species $j$ we need

$$\frac{1}{N_i}\frac{dN_i}{dt}\Big|_{N_i \approx 0, \tilde{N}_j} \approx \left(r_i - (\gamma + m_i)\right) \left(1 - \frac{r_i\alpha_i}{r_i - (\gamma + m_i)}\, 0 - \frac{r_i\alpha_i}{r_i - (\gamma + m_i)} \tilde{N}_j\right) > 0.$$

Plugging in $\tilde{N}_j = \frac{r_j - (\gamma + m_j)}{r_j\alpha_j}$ (and writing out the similar condition for species $j$) yields the following conditions for species $i$ to invade $j$ (and $j$ to invade $i$ respectively):

$$\frac{r_i\alpha_i}{r_i - (\gamma + m_i)} < \frac{r_j\alpha_j}{r_j - (\gamma + m_j)}, \qquad \frac{r_i\alpha_i}{r_i - (\gamma + m_i)} > \frac{r_j\alpha_j}{r_j - (\gamma + m_j)}$$

which clearly cannot be satisfied simultaneously. (The conditions for a feasible coexistence equilibrium point in this model are similar.) So when the species densities stay low enough that there is no critical patch-age at which they block offspring survival entirely (the region in which these Lotka-Volterra equations apply), the two species cannot stably coexist from disturbance and patch dynamics.

If instead the linear density-dependent term goes to 0 at some critical patch-age $a_{c,i}$, then, by enforcing the max function and changing the upper limit of integration, we have

$$\frac{dN_i}{dt} = r_i N_i \int_0^{a_{c,i}} da\ \left(1 - \alpha_i n_i(a,t) - \alpha_i n_j(a,t)\right) \rho(a,t) - (\gamma + m_i) N_i.$$

This would not necessarily give us macroscopic equations equivalent to the Lotka Volterra form. We will consider coexistence behaviors in the presence of a critical patch-age using an invasion analysis (see Sec. 4.2). We find that the invasion conditions only require consideration of the critical patch-age and differ from what one gets from these Lotka-Volterra equations if the species differ in the robustness of their offspring survival to competition, i.e. if they differ in their competition sensitivities: $\alpha_i \neq \alpha_j$.

Hence our competition acting on offspring-survival model case does not permit coexistence *unless* species vary their competition sensitivities $\alpha_i \neq \alpha_j$, and in particular will not permit coexistence simple from a reproduction/adult-survival trade-off as our other model cases do.

#### 4.2 Invasion Analysis with a Critical Patch-Age

Here we carry out an invasion analysis to predict coexistence when the assumptions leading to the average density Lotka-Volterra equations derived in the prior section (Sec. 4.1) *do not apply*, i.e. when the species densities *may* drive the linear density-dependent factor in the recruitment term to 0. In more biological terms, we here consider coexistence in this model case for instances where species may not be able to recruit into old patches at all, specifically when their offspring survival can be driven to zero by competition with adults already established in those patches.

##### 4.2.1 Single Species Equilibrium Density over Patch-Ages

The first step in our invasion analysis is to derive the equilibrium density of a species when on its own. We first consider the single species version of our dynamical equation (S18):

$$\frac{\partial n_i(a,t)}{\partial t} = r_i N_i(t) \max[0, (1 - \alpha_i n_i(a,t))] - m_i n_i(a,t) - \frac{\partial n_i(a,t)}{\partial a}.$$

If we define $a_{c,i}$ as the patch-age at which the linear density-dependent factor in the recruitment term goes to 0, then we know that for $a < a_{c,i}$ the equation can simply be written

$$\frac{\partial n_i(a,t)}{\partial t} = r_i N_i(t) \left(1 - \alpha_i n_i(a,t)\right) - m_i n_i(a,t) - \frac{\partial n_i(a,t)}{\partial a}.$$

Let's now denote the equilibrium dependence of species $i$ on patch-age when it is on its own as $\tilde{n}_{i|0}(a)$ (but dropping the explicit dependence on $a$ in intermediate steps below for the reader's convenience) and its average density as $\tilde{N}_{i|0}$. So we have, at equilibrium, (again for $a < a_{c,i}$)

$$0 = r_i \tilde{N}_{i|0}(1 - \alpha_i \tilde{n}_{i|0}) - m_i \tilde{n}_{i|0} - \frac{dn_i}{da}$$
$$0 = r_i \tilde{N}_{i|0} - (\alpha_i r_i \tilde{N}_{i|0} + m_i)\tilde{n}_{i|0} - \frac{dn_i}{da}$$

$$\int \frac{dn_i}{r_i \tilde{N}_{i|0} - (\alpha_i r_i \tilde{N}_{i|0} + m_i)\tilde{n}_{i|0}} = \int da$$
$$-\frac{1}{\alpha_i r_i \tilde{N}_{i|0} + m_i} \ln\left|r_i \tilde{N}_{i|0} - \left(\alpha_i r_i \tilde{N}_{i|0} + m_i\right)\tilde{n}_{i|0}\right| = a + C_0$$
$$r_i \tilde{N}_{i|0} - \left(\alpha_i r_i \tilde{N}_{i|0} + m_i\right)\tilde{n}_{i|0} = C_1 e^{-\left(\alpha_i r_i \tilde{N}_{i|0} + m_i\right)a}$$

resulting in

$$\tilde{n}_{i|0} = \frac{1}{\alpha_i r_i \tilde{N}_{i|0} + m_i}\left(r_i \tilde{N}_{i|0} - C_1 e^{-\left(\alpha_i r_i \tilde{N}_{i|0} + m_i\right)a}\right).$$

Using our boundary condition that the density of new patches is 0 (disturbance wipes patches clean), i.e. $n_i(0) = 0$ this results in

$$\tilde{n}_{i|0} = \frac{r_i \tilde{N}_{i|0}}{\alpha_i r_i \tilde{N}_{i|0} + m_i}\left(1 - e^{-\left(\alpha_i r_i \tilde{N}_{i|0} + m_i\right)a}\right) \qquad \text{for } a < a_{c,i}.$$

We could next consider the dynamics of $n_i(a)$ for $a < a_{c,i}$, but let's first further consider if when a single species is on its own its offspring survival would ever be driven to 0 in old patches, i.e. consider whether keeping track of a finite positive $a_{c,i}$ is even necessary. Our critical age can be solved for using

$$\begin{aligned} 0 &= 1 - \alpha_i n_i(a_{c,i}) \\ 0 &= 1 - \frac{\alpha_i r_i N_i}{\alpha_i r_i N_i + m_i}\left(1 - e^{-(\alpha_i r_i N_i + m_i)a_{c,i}}\right). \end{aligned}$$

Consider the case were $\lim_{a_{c,i}\to\infty}$. This case maximizes the term being subtracted from 1, and thus if it is not equal to 0, a smaller $a_{c,i}$ would not solve this equation either.

$$\begin{aligned} 0 &= \lim_{a_{c,i}\to\infty}\left(1 - \frac{\alpha_i r_i \tilde{N}_{i|0}}{\alpha_i r_i \tilde{N}_{i|0} + m_i}\left(1 - e^{-\left(\alpha_i r_i \tilde{N}_{i|0} + m_i\right)a_{c,i}}\right)\right) \\ 0 &= 1 - \frac{\alpha_i r_i \tilde{N}_{i|0}}{\alpha_i r_i \tilde{N}_{i|0} + m_i} \end{aligned}$$

Seeing as all our parameters are positive, we can clearly see that the term on the RHS will never equal 0, because the numerator is less than the denominator. Thus $a_{c,i}$ does <u>not</u> exist in the single species case.

Thus we have, for all $a$,

$$\tilde{N}_{i|0} = \frac{r_i \tilde{N}_{i|0}}{\alpha_i r_i \tilde{N}_{i|0} + m_i}\left(1 - e^{-\left(\alpha_i r_i \tilde{N}_{i|0} + m_i\right)a}\right).$$

Using the definition of the total density at equilibrium of one species on its own, $\tilde{N}_{i|0}$, we can derive an expression for it in terms of model parameters. In other words, using

$$\tilde{N}_{i|0} = \int_0^\infty \rho(a)\tilde{n}_{i|0}(a)da$$

we can plug in our expression for $\tilde{n}_{i|0}(a)$ in terms of $\tilde{N}_{i|0}$ and then solve:

$$\begin{aligned} \tilde{N}_{i|0} &= \int_0^\infty \gamma e^{-\gamma a}\frac{r_i \tilde{N}_{i|0}}{\alpha_i r_i \tilde{N}_{i|0} + m_i}\left(1 - e^{-\left(\alpha_i r_i \tilde{N}_{i|0} + m_i\right)a}\right) da \\ \tilde{N}_{i|0} &= \frac{\gamma r_i \tilde{N}_{i|0}}{\alpha_i r_i \tilde{N}_{i|0} + m_i}\int_0^\infty e^{-\gamma a}\left(1 - e^{-\left(\alpha_i r_i \tilde{N}_{i|0} + m_i\right)a}\right) da \\ \tilde{N}_{i|0} &= \frac{\gamma r_i \tilde{N}_{i|0}}{\alpha_i r_i \tilde{N}_{i|0} + m_i}\left(\frac{\alpha_i r_i \tilde{N}_{i|0} + m_i}{\gamma\left(\alpha_i r_i \tilde{N}_{i|0} + \gamma + m_i\right)}\right) \\ \tilde{N}_{i|0} &= \frac{r_i \tilde{N}_{i|0}}{\alpha_i r_i \tilde{N}_{i|0} + \gamma + m_i}. \end{aligned}$$

Finally we have

$$\tilde{N}_{i|0} = \frac{r_i - (m_i + \gamma)}{r_i \alpha_i}.$$

This results in the one species density at equilibrium

$$\tilde{n}_{i|0} = \frac{1}{\alpha_i}\left(1 - \frac{m_i}{(r_i - \gamma)}\right)\left(1 - e^{-a(r_i - \gamma)}\right). \tag{S19}$$

##### 4.2.2 Solving for Critical Patch-Age and Carrying out Invasion Analysis

Now we use this expression ((S19)) for the equilibrium of a single species to ask if species $i$ can invade a population of species $j$ that is at equilibrium on its own. We can, using an approach entirely analogous to the one we used for the competition acting on reproduction (see Sec. 3.4), derive an expression for the invasion growth rate of the average density of the species analogous to (S8). For this model case we get

$$\frac{1}{N_i}\frac{dN_i}{dt}\Big|_{\left(\epsilon_i(a), \tilde{n}_{j|0}(a)\right)} = \frac{r_i \int_0^\infty \gamma e^{-\gamma a}\epsilon_i(a)da \int_0^\infty \rho(a)\max\left[0, \left(1 - \alpha_i \tilde{n}_{j|0}(a)\right)\right]}{\int_0^\infty \gamma e^{-\gamma a}\epsilon_i(a)da} - (m_i + \gamma).$$

We first note that the integral over $\epsilon_i(a)$ in this expression simply cancels out. So this expression simplifies to

$$\frac{1}{N_i}\frac{dN_i}{dt}\Big|_{\left(\epsilon_i(a),\tilde{n}_{j|0}(a)\right)} = r_i \int_0^{\infty} \rho(a) \max\left[0, \left(1-\alpha_i \tilde{n}_{j|0}(a)\right)\right] - (m_i+\gamma)\,.$$

Now we consider at what patch-age the invading species $i$ would have its linear density-dependent term go to 0, so that we can implement the max function by simply putting a finite limit on the integral. In other words we will calculate this invasion growth rate as

$$\frac{1}{N_i}\frac{dN_i}{dt}\Big|_{\left(\epsilon_i(a),\tilde{n}_{j|0}(a)\right)} = r_i \int_0^{a_{c,j|0}} \rho(a) \left(1-\alpha_i \tilde{n}_{j|0}(a)\right) - (m_i+\gamma)\,.$$

Let's call this critical patch-age $a_{ci,j|0}$. To solve for it we use the following

$$1-\alpha_i \tilde{n}_{j|0}(a_{ci,j|0}) = 0$$

$$1-\alpha_i \frac{1}{\alpha_j}\left(1-\frac{m_j}{(r_j-\gamma)}\right)\left(1-e^{-a_{ci,j|0}(r_j-\gamma)}\right) = 0.$$

Here we first note that for the actual $a_{ci,j|0}$ to be positive and finite we require that the value of the term being subtracted from 1 be greater than 1 when $a_{ci,j|0} \to \infty$, i.e. we need

$$\frac{\alpha_i}{\alpha_j}\left(1-\frac{m_j}{r_j-\gamma}\right) > 1.$$

This can be rewritten as

$$\frac{\alpha_i}{\alpha_j}\frac{r_j-(m_j+\gamma)}{r_j-\gamma} > 1.$$

Since the second fraction on the left hand side is $< 1$, this means that at a minimum we need

$$\alpha_i > \alpha_j$$

for there to be a finite positive $a_{ci,j|0}$, i.e. for species $j$ to fully block the offspring survival of species $i$ in older patches. Unless this blocking occurs, the invasion of species $i$ would be no different than what is predicted by the Lotka-Volterra model derived in the prior section (Sec. 4.1), and there will mutual invasion. So in what remains we assume $\alpha_i > \alpha_j$.
Next we solve for $a_{ci,j|0}$ using the equation above, and get

$$a_{ci,j|0} = \frac{1}{r_j-\gamma}\ln\left|\frac{\alpha_i(\gamma+m_j-r_j)}{\alpha_i m_j - (\alpha_i-\alpha_j)(r_j-\gamma)}\right|.$$

Carrying out the integration and substituting this integration limit results in the following expression for the invasion growth rate of species $i$ into an equilibrium population of species $j$.

$$\frac{1}{N_i}\frac{dN_i}{dt}\Big|_{\left(\epsilon_i(a),\tilde{n}_{j|0}(a)\right)} = r_i\left(1-\frac{\alpha_i}{\alpha_j}\frac{r_j-(\gamma+m_j)}{r_j}\right) - (\gamma+m_i) + r_i\frac{\alpha_i}{\alpha_j}\left(\frac{(\alpha_i-\alpha_j)(r_j-\gamma)-m_i\alpha_i}{\alpha_i(r_j-(m_j+\gamma))}\right)^{\frac{r_i}{r_i-\gamma}} \tag{S20}$$

For species $j$, which we have now assumed has $\alpha_j < \alpha_i$, we do not need to worry about the critical patch-age for its invasion growth rate. Species $i$'s equilibrium density will never be high enough to block species $j$ in old patches. So the Lotka-Volterra model invasion growth rate conditions can be used for it to consider mutual invasion. Alternatively, we can follow the steps analogous to the above and get the same statement:

$$\begin{aligned}\frac{1}{N_j}\frac{dN_j}{dt}\Big|_{\left(\tilde{n}_{i|0}(a),\epsilon_j(a)\right)} &= \frac{r_j\int_0^{\infty}\rho(a)\epsilon_j(a)\left(1-\alpha_j\tilde{n}_{i|0}(a)\right) - (m_j+\gamma)}{\int_0^{\infty}\rho(a)\epsilon_j(a)da} \\ &= r_j\left(1-\frac{\alpha_j}{\alpha_i}\frac{r_i-(\gamma+m_i)}{r_i}\right) - (\gamma+m_j)\,. \end{aligned} \tag{S21}$$

Setting this $> 0$ we can show it is equivalent to the condition for species $j$ to invade species $i$ from the Lotka-Volterra model in the prior section (Sec. 4.1). We can also see that the invasion condition for species $i$ is equal to what it would be predicted to be from the Lotka-Volterra equations in the prior section (in which case it will be negative when the invasion growth rate of $j$ is positive) plus a term that can be positive. So we can see right away a potential for coexistence where there was none in the Lotka-Volterra model.

We used numerical evaluation to consider whether these two invasion conditions overlap and predict mutual invasion, and then validated the coexistence regions we found with numerical simulations. We found that a trade-off between reproduction and robustness of offspring-survival to competition (i.e. $r_i > r_j$ and $\alpha_i < \alpha_j$) could produce coexistence while a trade-off between adult-survival and robustness of offspring-survival does not (i.e. $m_i < m_j$ and $\alpha_i < \alpha_j$). We found that there was some limit to the similarity of coexistence of two species however–we could not find coexistence with species $i$ arbitrarily similar to species $j$ in its demographic parameters–the coexistence cone started at a point some away from the demographic parameters of species $j$. See Fig. 5 for a comparison of analytical and numerical coexistence regions and Fig. 6 for a look at how varying disturbance alters the coexistence region.

### 5 Competition Acting on Adult-Survival

The dynamical equation for a species' density over patch-age in this case, with a general form $f_i()$ for the density-dependence of per-capita mortality, is

$$\frac{\partial n_i(a,t)}{\partial t} = \underbrace{r_i \int_0^\infty \rho(a',t) n_i(a',t)\, da'}_{\text{recruitment}} - \underbrace{m_i f_i(n_i(a,t), n_j(a,t)) n_i(a,t)}_{\text{mortality}} - \underbrace{\frac{\partial n_i}{\partial a}}_{\substack{\text{patch-}\\\text{aging}}}$$

Assuming the per-capita mortality rate increases linearly with density we have

$$\frac{\partial n_i(a,t)}{\partial t} = \underbrace{r_i \int_0^\infty \rho(a',t) n_i(a',t) da'}_{N_i(t)} - m_i \left(1 + n_i(a,t) + n_j(a,t)\right) n_i(a,t) - \frac{\partial n_i}{\partial a}$$

$$\begin{aligned} \frac{\partial \rho(a,t)}{\partial t} &= -\frac{\partial \rho(a,t)}{\partial a} - \gamma \rho(a,t) \\ \rho(0,t) &= \int_0^\infty \gamma \rho(a,t) da \\ n_i(0,t) &= 0. \end{aligned}$$

This partial differential question is different from our other two model cases in that it has a *nonlinear* dependence on the density of the focal species $i$ in the focal patch of age $a$. As a result, it was more difficult to approach analytically, though we were able to make some minimal progress in the single species case.

Specifically, we were able to show that the density of species $i$ over patch-ages at equilibrium ($\tilde{n}_i$) will follow

$$\tilde{n}_i(a) = \frac{2 r_i \tilde{N}_i}{\sqrt{m_i \left(4 r_i \tilde{N}_i + m_i\right)} \coth\left(\frac{1}{2} a \sqrt{m_i \left(4 r_i \tilde{N}_i + m_i\right)}\right) + m_i} \tag{S22}$$

where $\tilde{N}_i$ is the average density across all patches at equilibrium. This form for the equilibrium density is plotted in Fig. 7.

However, we were unable to solve for the average density $\tilde{N}_i$ at equilibrium in terms of model parameters, as the necessary integrals yielded only Hypergeometric functions:

$$\begin{aligned} N_i(a) &= \int da\; n_i(a) \rho(a) \\ &= \int_0^\infty da\; \gamma e^{-\gamma a} \frac{2 r_i N_i \left(e^{a\sqrt{m_i(4 r_i N_i + m_i)}} - 1\right)}{\sqrt{m_i \left(4 r_i N_i + m_i\right)} \left(e^{a\sqrt{m_i(4 r_i N_i + m_i)}} + 1\right) + m_i \left(e^{a\sqrt{m_i(4 r_i N_i + m_i)}} - 1\right)} \\ &= 2 r_i N_i \gamma \int_0^\infty da\; e^{-\gamma a} \frac{\left(e^{a\sqrt{m_i(4 r_i N_i + m_i)}} - 1\right)}{\sqrt{m_i \left(4 r_i N_i + m_i\right)} \left(e^{a\sqrt{m_i(4 r_i N_i + m_i)}} + 1\right) + m_i \left(e^{a\sqrt{m_i(4 r_i N_i + m_i)}} - 1\right)}. \end{aligned}$$

Integrating this results in

$$\begin{aligned} &\frac{2 r_i N_i}{\left(\sqrt{m_i \left(4 r_i N_i + m_i\right)} + m_i\right) \left(\sqrt{m_i \left(4 r_i N_i + m_i\right)} + \gamma\right)} \\ &\left(\left(\sqrt{m_i \left(4 r_i N_i + m_i\right)} + \gamma\right) \text{Hypergeometric2F1}\left[1, \frac{\gamma}{\sqrt{m_i \left(4 r_i N_i + m_i\right)}}, \frac{\gamma}{\sqrt{m_i \left(4 r_i N_i + m_i\right)}} + 1, \frac{m_i - \sqrt{m_i \left(4 r_i N_i + m_i\right)}}{\sqrt{m_i \left(4 r_i N_i + m_i\right)} + m_i}\right]\right. \\ &\quad \left. - \gamma\, \text{Hypergeometric2F1}\left[1, \frac{\gamma}{\sqrt{m_i \left(4 r_i N_i + m_i\right)}} + 1, \frac{\gamma}{\sqrt{m_i \left(4 r_i N_i + m_i\right)}} + 2, \frac{m_i - \sqrt{m_i \left(4 r_i N_i + m_i\right)}}{\sqrt{m_i \left(4 r_i N_i + m_i\right)} + m_i}\right]\right) \end{aligned}$$

Hence we were unable to carry out an analytical invasion analysis in terms of model parameters for this model case. Instead we approached this model using our numerical simulation method, carrying out mutual invasion analyses over the parameter space that would include species variation along hypothesized demographic trade-offs.

We did indeed find coexistence when species varied along both a reproduction versus adult-survival trade-off in (See Fig. (8), and along a trade-off between reproduction and robustness of survival to competition (See Fig. 9).

### 6 Numerical Method

There are many numerical methods for simulating structured population models. Common methods include the Escalator Boxcar Train (EBT), fixed mesh finite volume, and moving mesh schemes. These methods each have their respective benefits and disadvantages which depend on the problem at hand (see [46] for a description and comparison). EBT was designed to avoid numerical diffusion that can occur with other schemes. However, the EBT method is of 1$^{\text{st}}$ order accuracy and therefore is slower (a finer discretization is required to achieve accuracy comparable to higher-order methods and this takes more computation time). Since the model we present is age-structured, we don't need to be as concerned about numerical diffusion, as it is exacerbated by growth rate related complexity. Therefore, we can prioritize speed of the simulation. To this end, we opt for a more accurate (and hence faster) $2^{\text{nd}}$ order scheme, specifically a finite volume scheme which makes use of a flux limiter method in the event of any sharp changes in the population density to avoid numerical diffusion. Specifically, we use a minmod flux limiter scheme which, heuristically, interpolates between high order and low order approximations of the flux. The weight of this interpolation is chosen based on a comparison of the forward and backward derivative at each point. By doing so, the scheme combats numerical artifacts commonly associated with higher order schemes, e.g., numerical diffusion, dissipation, and over-shooting of local extrema [47], while preserving the high order convergence of the scheme globally (in an $L^1$ sense).

We now describe the numerical methods used in the paper. For given positive mesh sizes $\Delta t$ and $\Delta a$, we discretize the patch-age domain uniformly by cells $\Lambda_j$ given by

$$\Lambda_j := \left[a_j - \frac{\Delta a}{2}, a_j + \frac{\Delta a}{2}\right],$$

where $a_j := j\Delta a$ for $j = 1, 2, \ldots, J$ (with the left bound modified for $j = 0$ to avoid negative ages). Our numerical scheme was designed to approximate the discretized solution of our general structured model framework equation at time $t_k := k\Delta t$. In other words, we calculate the values $n_j^k$ and $\rho_j^k$ which approximate the exact solution in the following way:

$$n_j^k \approx \frac{1}{|\Lambda_j|}\int_{\Lambda_j} n(t_k, a)da \quad \text{and} \quad \rho_j^k \approx \frac{1}{|\Lambda_j|}\int_{\Lambda_j} \rho(t_k, a)da.$$

We highlight the defining property of a flux limiter method by considering the continuous time evolution equation for $\rho$ as an example and letting $\bar{\rho}_j(t) := \frac{1}{|\Lambda_j|}\int_{\Lambda_j} \rho(t, a)da$. The evolution of the cell averages over time can be formulated by integrating the PDE over the cell to arrive at

$$\partial_t \bar{\rho}_j(t) = -\frac{1}{|\Lambda_j|}\int_{\Lambda_j} \partial_a \rho(t, a)da - \frac{1}{|\Lambda_j|}\int_{\Lambda_j} \gamma\rho(t, a)da = -\frac{1}{|\Lambda_j|}[\rho(t, a_j + \frac{\Delta a}{2}) - \rho(t, a_j - \frac{\Delta a}{2})] - \gamma\bar{\rho}_j(t). \quad \text{(S23)}$$

(S23) is the main branching point for higher-order finite volume schemes. The main question is how to best approximate the values $\rho(t, a_j + \frac{\Delta a}{2})$ and $\rho(t, a_j - \frac{\Delta a}{2})$ given the average values of the neighboring cells. Flux limiter methods approximate these values in a way which mitigate the aforementioned numerical artifacts. In particular, we make use of a minmod flux limiter which is a total variation diminishing (TVD) method, i.e., a method which ensures a decrease in oscillations over time. For more information on the general theory of flux limiter methods, we direct the reader to [47].

Using the notation above, we next describe the numerical scheme we use for all models in this paper. To avoid the over use of subscripts, we write the scheme as if only a single species ($n$) were present, but to emphasise the dependence on all present species we use the notation $\vec{n}$. We consider the numerical scheme:

$$\begin{cases} \rho_j^{k+1} = \rho_j^k - \dfrac{\Delta t}{\Delta a}(f[\rho_j^k] - f[\rho_{j-1}^k]) - \Delta t \gamma_j^k \rho_j^k, & j = 1, 2, \ldots, J \\ \rho_0^k = \Delta a \displaystyle\sum_{j=1}^{J}{}^{*} \rho_j^k \gamma_j^k & \\ n_j^{k+1} = n_j^k + \Delta t\, r_j^k(\vec{n}^k, \rho^k) - \dfrac{\Delta t}{\Delta a}(f[n_j^k] - f[n_{j-1}^k]) - \Delta t\, d_j^k n_j^k, & j = 1, 2, \ldots, J \\ n_0^k = 0 & \end{cases}$$

where the flux term is given by

$$f[m_j^k] := \begin{cases} m_j^k + \frac{1}{2}\,\mathrm{mm}(m_{j+1}^k - m_j^k, m_j^k - m_{j-1}^k) & j = 2, 3, \ldots, J-2 \\ m_j^k & j = 0, 1, J-1, J \end{cases}$$

and the sum on the boundary is given by

$$\sum_{j=1}^{J}{}^{*} \rho_j^k \gamma_j^k := \frac{3}{2}\rho_1^k \gamma_1^k + \frac{1}{2}\rho_J^k \gamma_J^k + \sum_{j=2}^{J-1} \rho_j^k \gamma_j^k.$$

Here we denote by $\mathrm{mm}(a, b)$ the minmod function given by

$$\mathrm{mm}(a, b) = \frac{\mathrm{sgn}(a) + \mathrm{sgn}(b)}{2}\min(|a|, |b|).$$

Relating back to the intuition from (S23), we see the numerical flux, $f[\rho_j^k]$, approximates the value $\rho(t_k, a_j + \frac{\Delta a}{2})$ in a similar manner to the Taylor expansion:

$$\rho(t_k, a_j + \frac{\Delta a}{2}) \approx \rho(t_k, a_j) + \frac{\Delta a}{2}\partial_a \rho(t_k, a_j).$$

When the signs of the forwards and backwards difference agree, the minmod function approximates the derivative, $\partial_a \rho(t_k, a_j)$, using the difference corresponding to the smaller magnitude. This helps to alleviate numerical diffusion around discontinuities by limiting the flux of mass leaving the cell $\Lambda_j$ and traveling into the neighboring cells. On the other hand, when the forwards and backwards differences have different signs, we expect a local extrema to be in the cell $\Lambda_j$. Therefore, the minmod function returns the value 0 and the flux reduces to the low resolution approximation $\rho(t_k, a_j + \frac{\Delta a}{2}) \approx \bar{\rho}_j(t_k)$ to avoid the creation of artificial extrema.

The approximation of the reproduction function, $r$, will vary depending on its form, but to maintain the expected second order convergence rate, one should use a second order method or better. For instance, for the case of density-independent recruitment, a third order quadrature method is used to calculate $r_j^k$. We remark that the above scheme is fully explicit. In other words, the calculation of information at any particular time step, depends exclusively on previously calculated information. As is common with explicit schemes, we require a stability condition on our mesh size. We take a course Courant–Friedrichs–Lewy (CFL) condition:

$$\zeta(\Delta t + \frac{3\Delta t}{2\Delta a}) \leq 1,$$

where $\zeta > 0$ bounds all model ingredients. Finally, for improvement on the time accuracy, we make use of the second order Runge-Kutta time discretization [48]. For more information on the stability and convergence of this method, we direct the reader to any of the following [47, 45, 42, 49, 50].

### List of Figures

1 **Comparing Analytical and Numerical Coexistence Regions with Competition Acting on Reproduction looking at a Reproduction/Adult-Survival Trade-Off.** Here we take a look at the numerical coexistence region (blue boxes) which overlay onto the analytical coexistence region (black; from (S16).) We have approximated where the two species critical patch-age cut-off value (green region) does not exist. We can see the analytical coexistence region is in the region where the critical patch-age cut-off value condition is satisfied. All figures have $\alpha_i = \alpha_j = 1$. . . . . . 22

2 **Analytical Coexistence Regions across Disturbance Rates with Competition Acting on Reproduction when looking at a Reproduction/Adult-Survival Trade-Off.** Intermediate levels of disturbance give us the widest coexistence region. When disturbance is too low, the species which depends on recently cleared patches is unable to gain a foothold on the landscape. When disturbance is too high, the species which is a better competitor and depends on older patches cannot survive. We begin graphing at the intersection of the black dashed lines where the mortality and reproduction of both species are equal: $m_i = m_j = 1, r_i = r_j = 5.6$. We consider the region where species $i$ has both a higher reproduction ($r_i$) and mortality ($m_i$) rate than species $j$ does. The blue region comes from (S6), the red from (S7) and the coexistence region comes from (S16). . . . . . 23

3 **Comparing Analytical and Numerical Coexistence Regions with Competition Acting on Reproduction looking at a Reproduction-Competition-Robustness/Adult-Survival Trade-Off.** Here we compare theoretical (black) and numerical (teal) coexistence regions, as well as the region verified numerically as having species' densities low enough that reproduction is not driven to zero at a finite patch-age due to competition (light green). We note that the numerical coexistence region (teal) is within the region where the numerical two-species coexistence condition is satisfied, and moved closer and closer to the analytical region as we refined the step sizes of our simulations. Note that our numerical simulations were only run over a limited range of parameters due to long run times. More accuracy requires a more refined step-size. The blue region comes from (S6), the red from (S7) and the coexistence region comes from (S16). Parameters are $r_j = r_i = 33.3$, $m_j = 18.5$, $\alpha_j = 7.6$ and $\gamma = 2.5$. . . . . . 24

4 **Comparing Analytical Coexistence Regions across Disturbance Rates with Competition Acting on Reproduction when looking at a Competition-Robustness/Adult-Survival Trade-Off.** Intermediate levels of disturbance give us the widest coexistence region. When disturbance is too low, the species which depends on recently cleared patches is unable to gain a foothold on the landscape. When disturbance is too high, the species which is a better competitor and depends on older patches cannot survive. Here we consider the region where species $i$ has a higher sensitivity to shading ($\alpha_i$) but a lower mortality ($m_i$) than species $j$. The blue region comes from (S6), the red from (S7) and the coexistence region comes from (S16). Our parameters here are $\mu_j = 18.5, \alpha_j = 7.6$, and $r_i = r_j = 33.3$. . . . . . 25

5 **Comparing Analytical and Numerical Coexistence Regions with Competition Acting on Offspring-Survival when looking at a Reproduction/Offspring-Survival Competition Robustness Trade-Off.** Here we compare our numerical (overlaid, teal boxes) and analytical results (red, blue and black regions) for the competition acting on offspring-survival case with parameters $m_i = m_j = 1, \alpha_j = 0.2, \gamma = 0.5$ and $r_j = 5$. The blue region comes from (S20), the red from (S21), and the coexistence region comes from the overlap of both of those. . . . . . 26

6 **Analytical Coexistence Regions across Disturbance Rates with Competition Acting on Offspring-Survival when looking at a Reproduction/Offspring-Survival Competition Robustness Trade-Off.** The white space extending from the cone to the coexistence region (black) here simply means that one species cannot invade the other–not that a species cannot persist. We can see how the coexistence region changes as we vary disturbance ($\gamma$). The blue region comes from (S20), the red from (S21), and the coexistence region comes from the overlap of both of those. Our parameters here are $m_i = m_j = 1, \alpha_j = 0.2$, and $r_j = 5$. . . . . . 27

#### Comparing Analytical and Numerical Coexistence Regions
##### Competition Acting on Reproduction

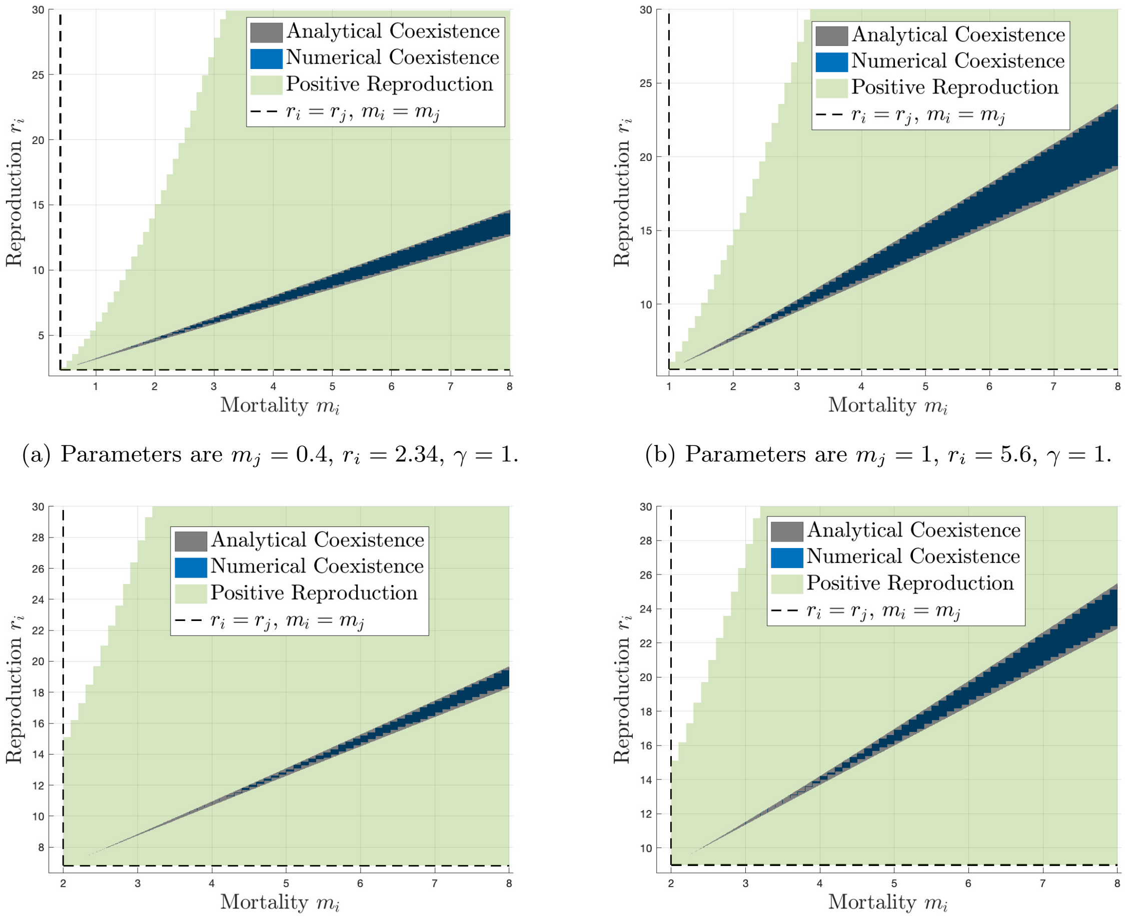

(a) Parameters are $m_j = 0.4$, $r_i = 2.34$, $\gamma = 1$.

(b) Parameters are $m_j = 1$, $r_i = 5.6$, $\gamma = 1$.

(c) Parameters are $m_j = 2$, $r_i = 6.8$, $\gamma = 1$.

(d) Parameters are $m_j = 2$, $r_i = 9$, $\gamma = 1$.

Figure 1: **Comparing Analytical and Numerical Coexistence Regions with Competition Acting on Reproduction looking at a Reproduction/Adult-Survival Trade-Off.** Here we take a look at the numerical coexistence region (blue boxes) which overlay onto the analytical coexistence region (black; from (S16).) We have approximated where the two species critical patch-age cut-off value (green region) does not exist. We can see the analytical coexistence region is in the region where the critical patch-age cut-off value condition is satisfied. All figures have $\alpha_i = \alpha_j = 1$.

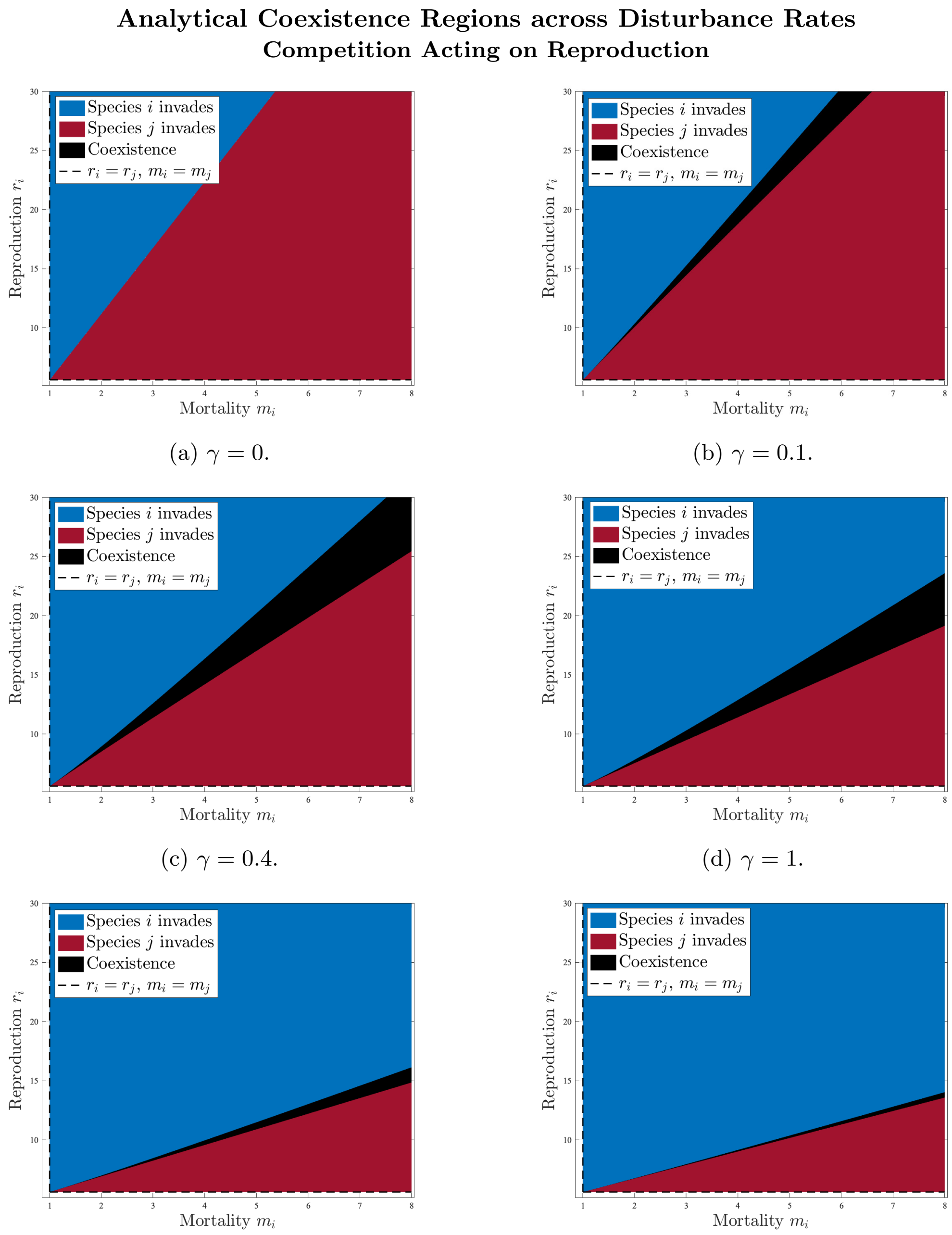

(a) $\gamma = 0$.

(b) $\gamma = 0.1$.

(c) $\gamma = 0.4$.

(d) $\gamma = 1$.

(e) $\gamma = 2.5$.

(f) $\gamma = 3.5$.

Figure 2: **Analytical Coexistence Regions across Disturbance Rates with Competition Acting on Reproduction when looking at a Reproduction/Adult-Survival Trade-Off.** Intermediate levels of disturbance give us the widest coexistence region. When disturbance is too low, the species which depends on recently cleared patches is unable to gain a foothold on the landscape. When disturbance is too high, the species which is a better competitor and depends on older patches cannot survive. We begin graphing at the intersection of the black dashed lines where the mortality and reproduction of both species are equal: $m_i = m_j = 1, r_i = r_j = 5.6$. We consider the region where species $i$ has both a higher reproduction ($r_i$) and mortality ($m_i$) rate than species $j$ does. The blue region comes from (S6), the red from (S7) and the coexistence region comes from (S16).

#### Comparing Analytical and Numerical Coexistence Regions
##### Competition Acting on Reproduction

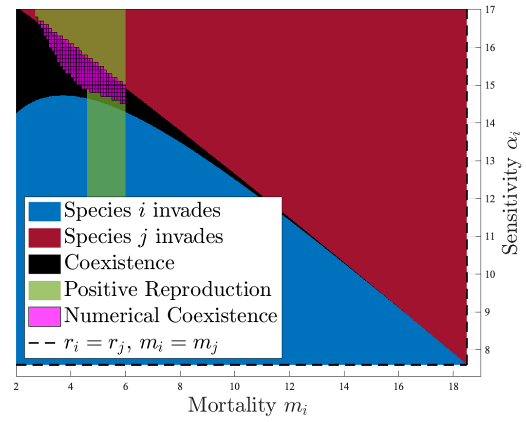

Figure 3: **Comparing Analytical and Numerical Coexistence Regions with Competition Acting on Reproduction looking at a Reproduction-Competition-Robustness/Adult-Survival Trade-Off.** Here we compare theoretical (black) and numerical (teal) coexistence regions, as well as the region verified numerically as having species' densities low enough that reproduction is not driven to zero at a finite patch-age due to competition (light green). We note that the numerical coexistence region (teal) is within the region where the numerical two-species coexistence condition is satisfied, and moved closer and closer to the analytical region as we refined the step sizes of our simulations. Note that our numerical simulations were only run over a limited range of parameters due to long run times. More accuracy requires a more refined step-size. The blue region comes from (S6), the red from (S7) and the coexistence region comes from (S16). Parameters are $r_j = r_i = 33.3$, $m_j = 18.5$, $\alpha_j = 7.6$ and $\gamma = 2.5$.

**Comparing Analytical Coexistence Regions across Disturbance Rates**
**Competition Acting on Reproduction**

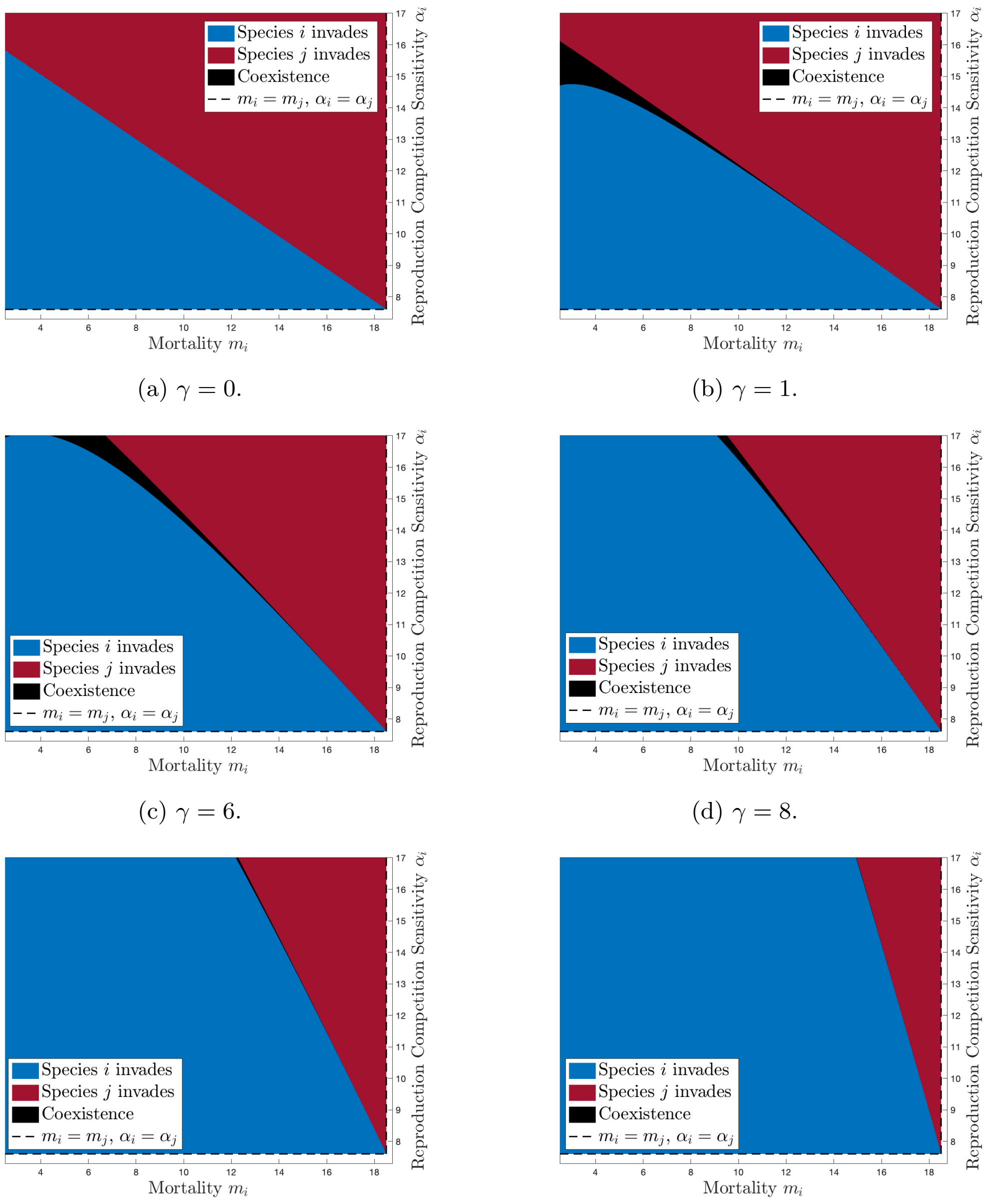

(a) $\gamma = 0$.

(b) $\gamma = 1$.

(c) $\gamma = 6$.

(d) $\gamma = 8$.

(e) $\gamma = 10$.

(f) $\gamma = 12$.

Figure 4: **Comparing Analytical Coexistence Regions across Disturbance Rates with Competition Acting on Reproduction when looking at a Competition-Robustness/Adult-Survival Trade-Off.** Intermediate levels of disturbance give us the widest coexistence region. When disturbance is too low, the species which depends on recently cleared patches is unable to gain a foothold on the landscape. When disturbance is too high, the species which is a better competitor and depends on older patches cannot survive. Here we consider the region where species $i$ has a higher sensitivity to shading ($\alpha_i$) but a lower mortality ($m_i$) than species $j$. The blue region comes from (S6), the red from (S7) and the coexistence region comes from (S16). Our parameters here are $\mu_j = 18.5, \alpha_j = 7.6$, and $r_i = r_j = 33.3$.

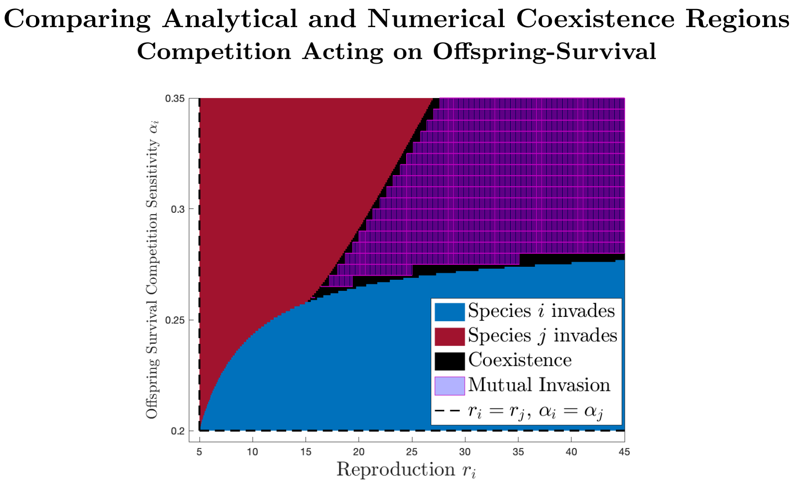

Figure 5: **Comparing Analytical and Numerical Coexistence Regions with Competition Acting on Offspring-Survival when looking at a Reproduction/Offspring-Survival Competition Robustness Trade-Off.** Here we compare our numerical (overlaid, teal boxes) and analytical results (red, blue and black regions) for the competition acting on offspring-survival case with parameters $m_i = m_j = 1, \alpha_j = 0.2, \gamma = 0.5$ and $r_j = 5$. The blue region comes from (S20), the red from (S21), and the coexistence region comes from the overlap of both of those.

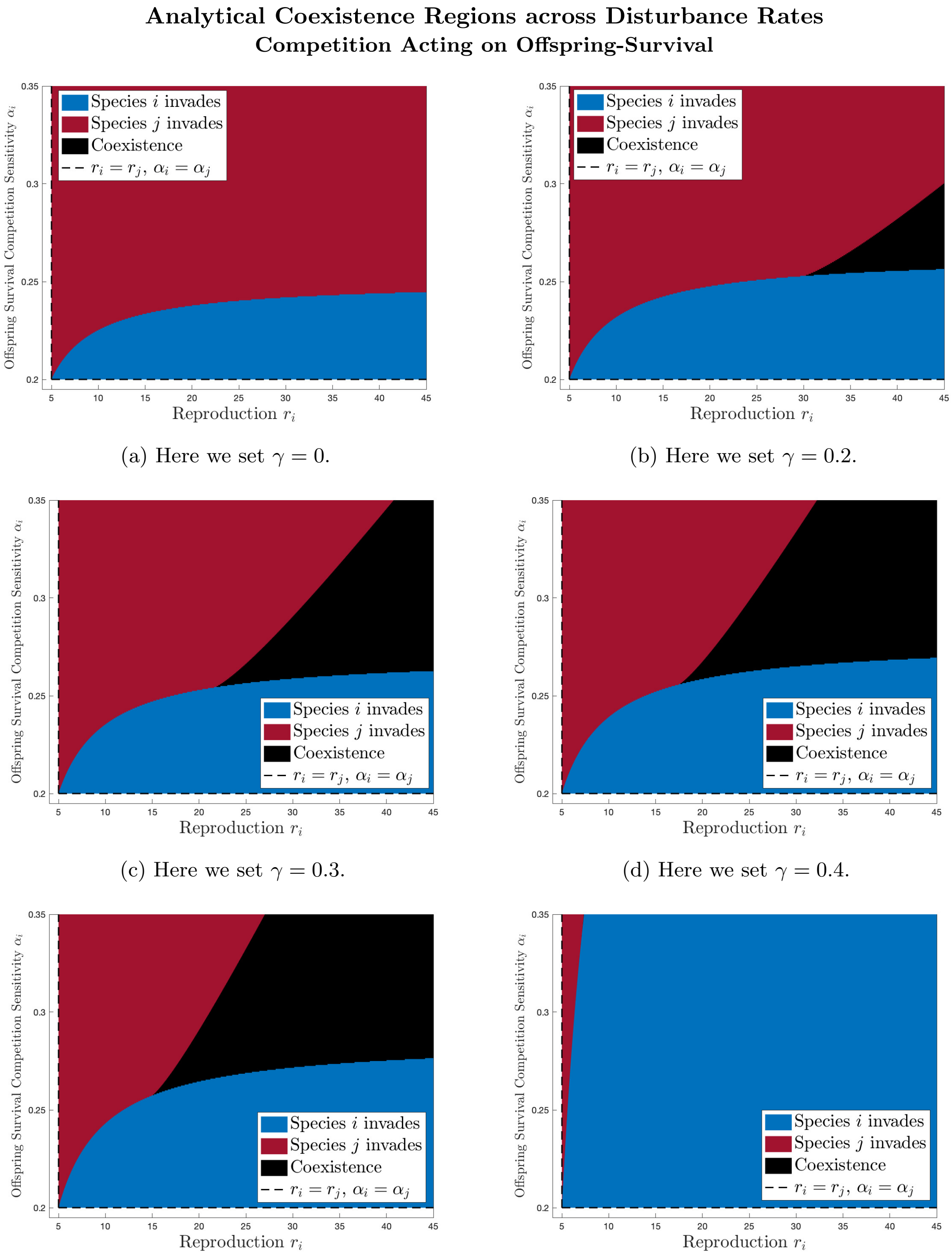

(a) Here we set $\gamma = 0$.

(b) Here we set $\gamma = 0.2$.

(c) Here we set $\gamma = 0.3$.

(d) Here we set $\gamma = 0.4$.

(e) Here we set $\gamma = 0.5$.

(f) Here we set $\gamma = 2.5$.

Figure 6: **Analytical Coexistence Regions across Disturbance Rates with Competition Acting on Offspring-Survival when looking at a Reproduction/Offspring-Survival Competition Robustness Trade-Off.** The white space extending from the cone to the coexistence region (black) here simply means that one species cannot invade the other–not that a species cannot persist. We can see how the coexistence region changes as we vary disturbance ($\gamma$). The blue region comes from (S20), the red from (S21), and the coexistence region comes from the overlap of both of those. Our parameters here are $m_i = m_j = 1, \alpha_j = 0.2$, and $r_j = 5$.

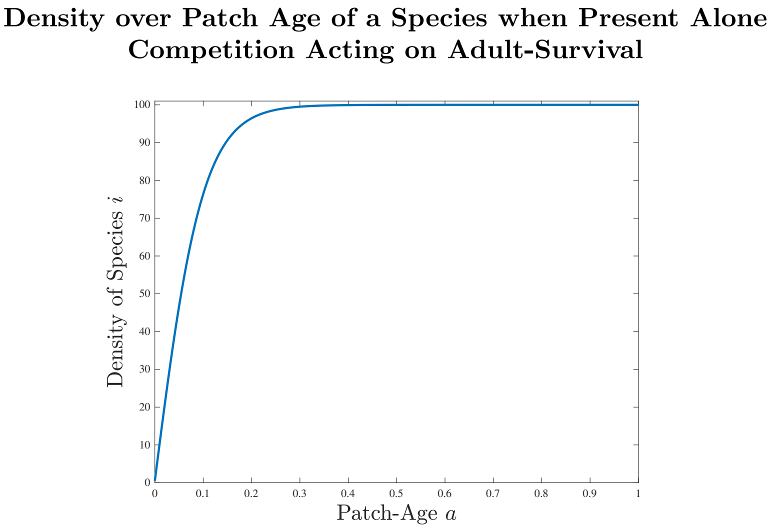

Figure 7: **Density over Patch Age of a Species when Present Alone with Competition Acting on Adult-Survival.** Graph of the density of species $i$ in the case of competition acting on adult-survival when $\tilde{N}_i = 100, r_i = 10, m_i = 0.1$. This graph comes from (S22).

**Comparing Numerical Coexistence Regions**
**Competition Acting on Adult-Survival**

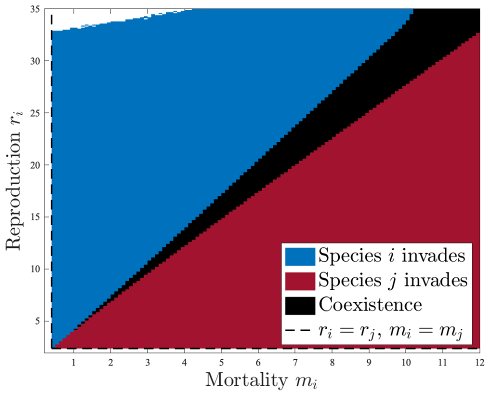

(a) Here we set $m_i = 0.4, r_i = 2.34, \gamma = 1$ and $\alpha = 1$.

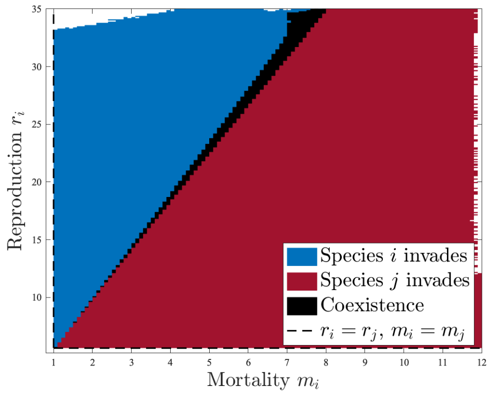

(b) Here we set $m_i = 1, r_i = 5.6, \gamma = 1$ and $\alpha = 1$.

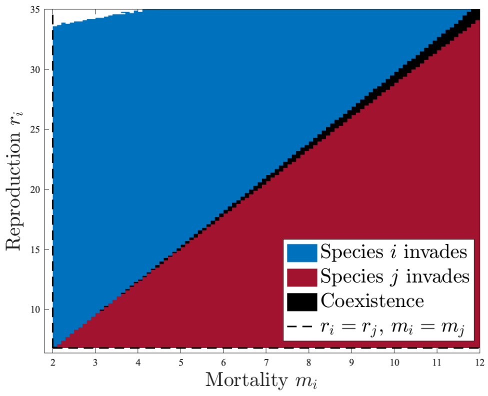

(c) Here we set $m_i = 2, r_i = 6.8, \gamma = 1$ and $\alpha = 1$.

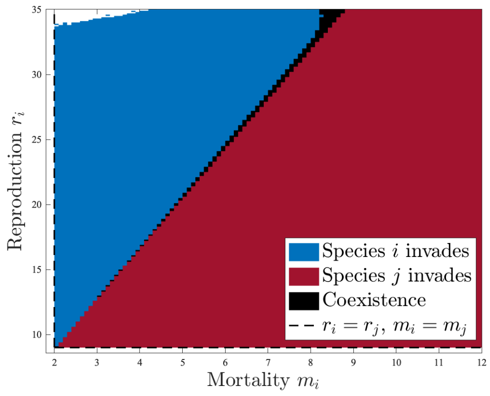

(d) Here we set $m_i = 2, r_i = 9, \gamma = 1$ and $\alpha = 1$.

Figure 8: **Comparing Numerical Coexistence Regions with Competition Acting on Adult-Survival when looking at a Reproduction/Adult-Survival Trade-Off.** The white space extending from the cone to the coexistence region (black) here simply means that one species cannot invade the other–not that a species cannot persist. The white edges near the top/sides of the graph are due to numerical inaccuracies.

**Comparing Coexistence Regions across Disturbance Rates**
**Competition Acting on Adult-Survival**

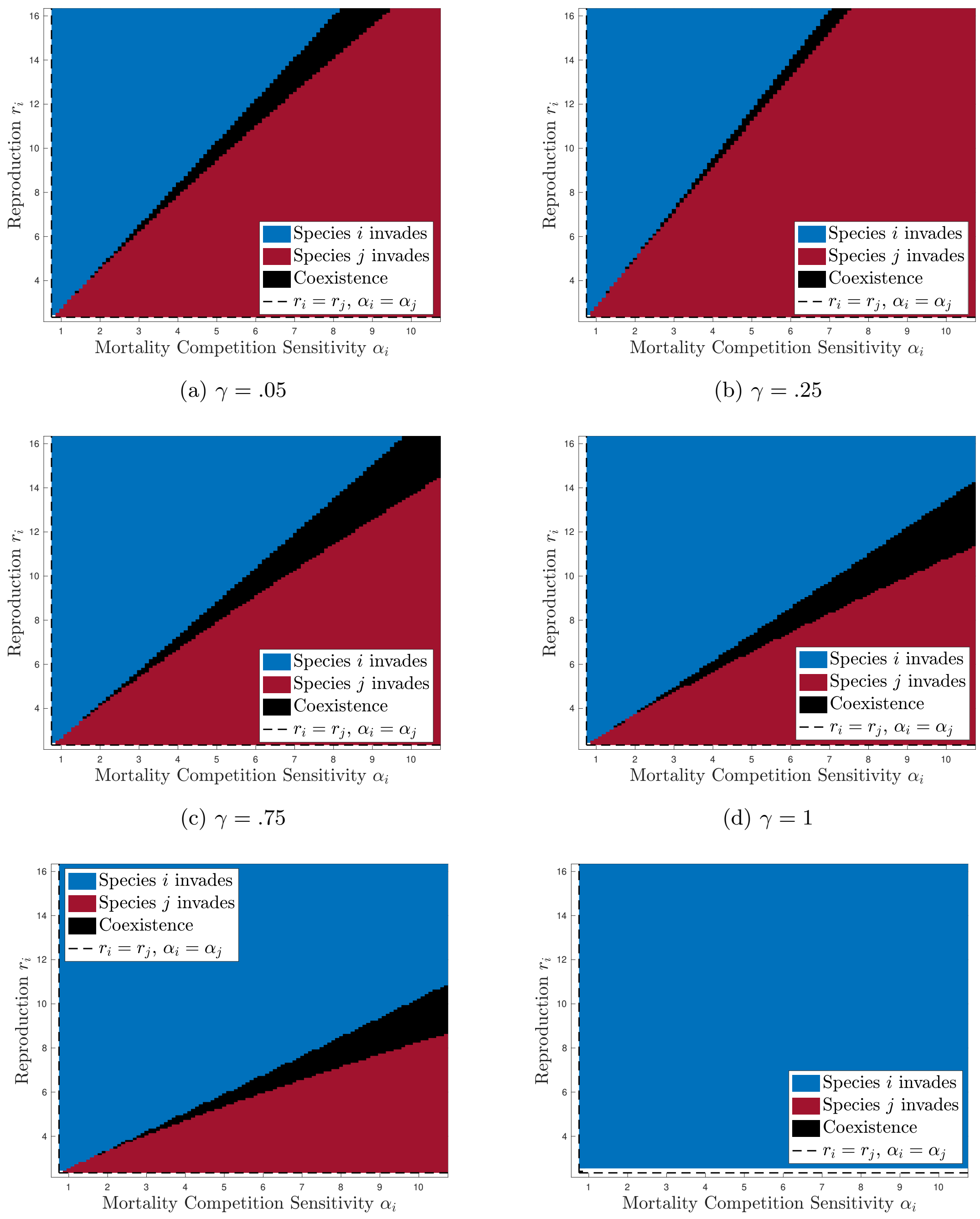

(a) $\gamma = .05$

(b) $\gamma = .25$

(c) $\gamma = .75$

(d) $\gamma = 1$

(e) $\gamma = 1.25$

(f) $\gamma = 2.$

Figure 9: **Comparing Coexistence Regions across Disturbance Rates with Competition Acting on Adult-Survival when looking at a Reproduction/Robustness-of-Adult-Survival-to-Competition Trade-Off.** Parameters are $m_j = m_i = 0.4$, $r_j = 2.34$, and $\alpha_j = 0.75$. Here we can see the coexistence region (black) changing as we increase the disturbance rate $\gamma$ from .05 to 2 while keeping the other parameters fixed.

### List of Tables

1 **Coexistence Results for Trade-Offs Considered.** For each model case we considered all demographic parameter trade-offs possible for that case for their potential to generate coexistence and whether or not coexistence is generated. The reproduction-adult survival trade-off was considered for all three model cases, and variation between species along that trade-off generated coexistence in all but the Competition acting on Offspring Survival case. The model cases consider in turn competition acting on one specific aspect of demography, and hence we can only consider robustness to competition of that component of demography competition is acting on for each case. We considered trade-offs between it and reproduction, and between it and adult survival, for each model case. Notable is that within-demographic component trade-offs, namely between reproduction and the robustness of reproduction to competition, and between survival and the robustness of survival to competition, did *not* enable coexistence. . . . . . . . . . . . . . . . . . . . . . . . . . . . . . . . . . . . . . . . . 32

| Competition on Reproduction | |
|---|---|
| Reproduction/Adult-Survival ($r_i/m_i$) | coexistence |
| Reproduction/Robustness of Reproduction to Competition ($r_i/\alpha_i$) | no coexistence |
| Robustness of Adult-Survival to Competition/Adult-Suvival ($\alpha_i/m_i$) | coexistence |

| Competition on Offspring-Survival | |
|---|---|
| Reproduction/Adult-Survival ($r_i/m_i$) | no coexistence |
| Reproduction/Robustness of Offspring-Survival to Competition ($r_i/\alpha_i$) | coexistence |
| Robustness of Adult-Survival to Competition/ | no coexistence |

| Competition on Adult-Survival | |
|---|---|
| Reproduction/Adult-Survival ($r_i/m_i$) | coexistence |
| Reproduction/Robustness of Adult-Survival to Competition ($r_i/\alpha_i$) | coexistence |
| Robustness of Adult-Survival to Competition/ Adult-Survival ($\alpha_i/m_i$) | no coexistence |

Table 1: **Coexistence Results for Trade-Offs Considered.** For each model case we considered all demographic parameter trade-offs possible for that case for their potential to generate coexistence and whether or not coexistence is generated. The reproduction-adult survival trade-off was considered for all three model cases, and variation between species along that trade-off generated coexistence in all but the Competition acting on Offspring Survival case. The model cases consider in turn competition acting on one specific aspect of demography, and hence we can only consider robustness to competition of that component of demography competition is acting on for each case. We considered trade-offs between it and reproduction, and between it and adult survival, for each model case. Notable is that within-demographic component trade-offs, namely between reproduction and the robustness of reproduction to competition, and between survival and the robustness of survival to competition, did *not* enable coexistence.
